# From labels to latents: revealing state-dependent computations and non-spatial replay with Jump Latent Variable Model

**DOI:** 10.64898/2025.12.14.694183

**Authors:** Zheyang (Sam) Zheng, Ipshita Zutshi, Roman Huszár, Yiyao Zhang, Mursel Karadas, György Buzsáki, Alex H. Williams

## Abstract

Neural activity is often interpreted by imposing external labels (e.g., position during locomotion) and decoding within that space (e.g. replay). While powerful, such supervision can mask structure in the data that do not correspond to the label. Unsupervised methods, in turn, often assume smooth latent dynamics and miss genuine discontinuities. We introduce a conceptually simple, computationally efficient latent variable model that infers both (i) the latent variables organizing population activity and (ii)whether their dynamics are continuous or fragmented in time. Fitting reduces to an expectation-maximization (EM) procedure that alternates two operations familiar to systems neuroscience—tuning-curve estimation and label decoding—without requiring external labels. Applied to rodent hippocampal recordings, our method identifies a continuity-fragmentation axis that characterizes population activity across sleep-wake brain states, modulated by cholinergic inputs. For a spatial navigation task, the model reveals distinct population patterns at the same physical position that supervised spatial decoding fails to detect. Moreover, the model enables replay analysis beyond spatial trajectories, and finds prevalent pre-play events associated with immobile and off-maze behaviors and post-task replay of locomotion. By not relying on externally imposed spatial labels, our approach exposes structure that supervised approaches obscure and provides a powerful tool for datasets lacking behavioral tracking.

## Introduction

Systems neuroscience commonly uses tuning curves and population decoding methods to quantify the information present in neural circuits (Dayan and Abbott, 2005; Pouget and Snyder, 2000; Averbeck et al., 2006; Quian Quiroga and Panzeri, 2009; Georgopoulos et al., 1986; Johnson et al., 2005; Angelaki et al., 2025; Brown et al., 1998; Hanks et al., 2015; Steinmetz et al., 2019; Meyers et al., 2008). Ironically, some of the most insightful results occure when the decoder prediction deviates from the ground truth label in structured ways, which we call “insight from mismatch”. One frequently studied type of mismatch is the “non-local representation” in the hippocampus. During running, the position of the animal decoded from hippocampal CA1 pyramidal neurons is often ahead or behind the animal’s actual position, revealing a non-veridical component of the spatial representation (Johnson and Redish, 2007; Gupta et al., 2012; Zheng et al., 2016; Kay et al., 2020; Johnson and Redish, 2007). During awake replay, the decoded position can also differ from the animal’s current position (Foster and Wilson, 2006; Diba and Buzsáki, 2007; Karlsson and Frank, 2009; Pfeiffer and Foster, 2013). In another example, Parks et al. (2024) used human consensus labels as ground truth, but discovered a more refined and faster time-scale sleep state transition where the label only indicates one state. These observations suggest there is structure in the population activity that is obscured or missed by the human-selected labels. Furthermore, in relating neural activities to labels that the experimenter deems important, neuroscientists often filter out data by semi-arbitrary criterion, for instance a speed threshold for analyzing spatial representation and replay in the hippocampus, while discarding most data during immobility. It could well be that by not restricting the analysis to preselected epochs (e.g. running), one can discover a less biased picture of what drives neuronal activity.

These limitations are pronounced in the study of episodic memory via replay. The predominant paradigm uses the spatial tuning curves of the hippocampal neurons during maze navigation as the template and decodes the spatial trajectories during sharp wave ripples (SPW-Rs) or population burst events (PBEs) during off-line periods (Davidson et al., 2009; Karlsson and Frank, 2009; Gupta et al., 2010; Dragoi and Tonegawa, 2011; Grosmark and Buzsáki, 2016; Farooq and Dragoi, 2019; Gillespie et al., 2021; Berners-Lee et al., 2022; Widloski and Foster, 2022; Liu et al., 2023; Mallory et al., 2025). While this approach has led to abundant discoveries, it begs the question whether only the spatial aspect of episodic memories is reflected in hippocampal replays, given the plethora of variables that the hippocampus has been found to encode (MacDonald et al., 2011, 2013; Aronov et al., 2017; Gauthier and Tank, 2018; Josselyn and Tonegawa, 2020; Zutshi et al., 2025).

These limitations motivate an unsupervised approach to population analysis and replay analysis. Dimensionality reduction and latent variable methods can leverage shared variability that organizes spiking activity to uncover low-dimensional trajectories and factors (Cunningham and Yu, 2014). While the dominant feature in the latent space is often explained by the variables that are known to be encoded in the region, the low-dimensional trajectories often exhibit additional visual features that prompt the inclusion of additional labels (Williams et al., 2018; Yang et al., 2024).

Existing unsupervised approaches often require smoothing of the spikes beforehand (e.g. before PCA or UMAP, Yang et al., 2024) or build in temporal smoothness in the latent trajectories (Yu et al., 2008; Wu et al., 2017; Pandarinath et al., 2018; George et al., 2024). However, neural activities routinely exhibit abrupt changes. For instance, hippocampal activities can represent remote locations during both the theta state and sharp wave ripples, abruptly deviating from the ongoing sequence (Johnson and Redish, 2007; Kay et al., 2020; Karlsson and Frank, 2009). When the animal enters sleep as well as transitioning between different stages of sleep, activities change abruptly in the brain (Saper et al., 2010). In the neocortex, spontaneous transitions between up-and down-state occur during sleep (Steriade et al., 1993). Sensory circuits traverse metastable attractors with sudden transitions during perceptual decision-making (Mazzucato et al., 2019; Miller and Katz, 2010). Prefrontal ensembles also reorganize discretely at rule or context shifts in working-memory tasks (Durstewitz et al., 2010). Moreover, the abrupt changes can last for extended periods of time, where the ensemble activities from time to time lose temporal autocorrelation, called a “fragmented” state, as seen during many hippocampal activities during sharp wave ripples (Denovellis et al., 2021, SPW-R). These findings motivate including “jumping” as a possibility for the evolution of latents.

We thus propose Jump Latent Variable Model (JumpLVM), an unsupervised method that extracts a low-dimensional latent variable in time, relates neural activities to the latent via latent tuning curves, and also infers the dynamics type of the latent variable, i.e. whether it evolves smoothly or in a fragmented way. It is a variant of the Gaussian Process Latent Variable Model (Lawrence, 2003; Wu et al., 2017; Jensen et al., 2020, GPLVM), with the possibility of non-smooth jump dynamics in the latent. In contrast to previous formulations that are mathematically complex, our inference is simple and efficient: we iteratively estimate tuning curves and apply Bayesian decoding—two routines already widely used by neuroscientists—to fit the model. After being fit to the data during the behavior epoch, the model extracts latent tuning curves in an unbiased manner. These templates can then be used for replay analysis during offline periods.

We demonstrate the utility of JumpLVM in extracellular recordings of hippocampal neural populations. We show that: 1) a continuity-fragmentation axis characterizes brain states across wake and sleep, co-varying with acetylcholine (ACh) levels; 2) unsupervised latents reveal distinct population patterns at the same physical position that supervised spatial decoding obscures; 3) these patterns are reactivated and replayed in distinct ways depending on the behavioral state during encoding.

## Results

### Iterative tuning-curve estimation and Bayesian decoding yield a flexible latent-variable model

A common analysis pipeline for neural population data is to (a) compute tuning curves with respect to some behavioral label, such as spatial position, and then (b) use Bayes rule to decode the behavioral label that is represented by a neural population (Zhang et al., 1998) (Fig.1A, purple arrows). In this paper, we investigate an approach where these two steps are iteratively repeated until convergence. That is, we treat the decoded representation (output of step b) as new labels for the next tuning curve update. These new tuning curves are then used to re-decode new labels, iterating until convergence. This procedure is equivalent to the expectation-maximization algorithm for a latent variable model (Dempster et al., 1977, EM), which discovers and organizes population activity patterns into a low-dimensional latent manifold (Fig. 1A, blue arrows). The yielded advantage is that we can discover patterns of neural coactivation without being biased by pre-selected labels (although human-supervised labels are often a useful way to initialize this unsupervised algorithm). Further details on the connection to EM are given in Methods.

**Figure 1.**
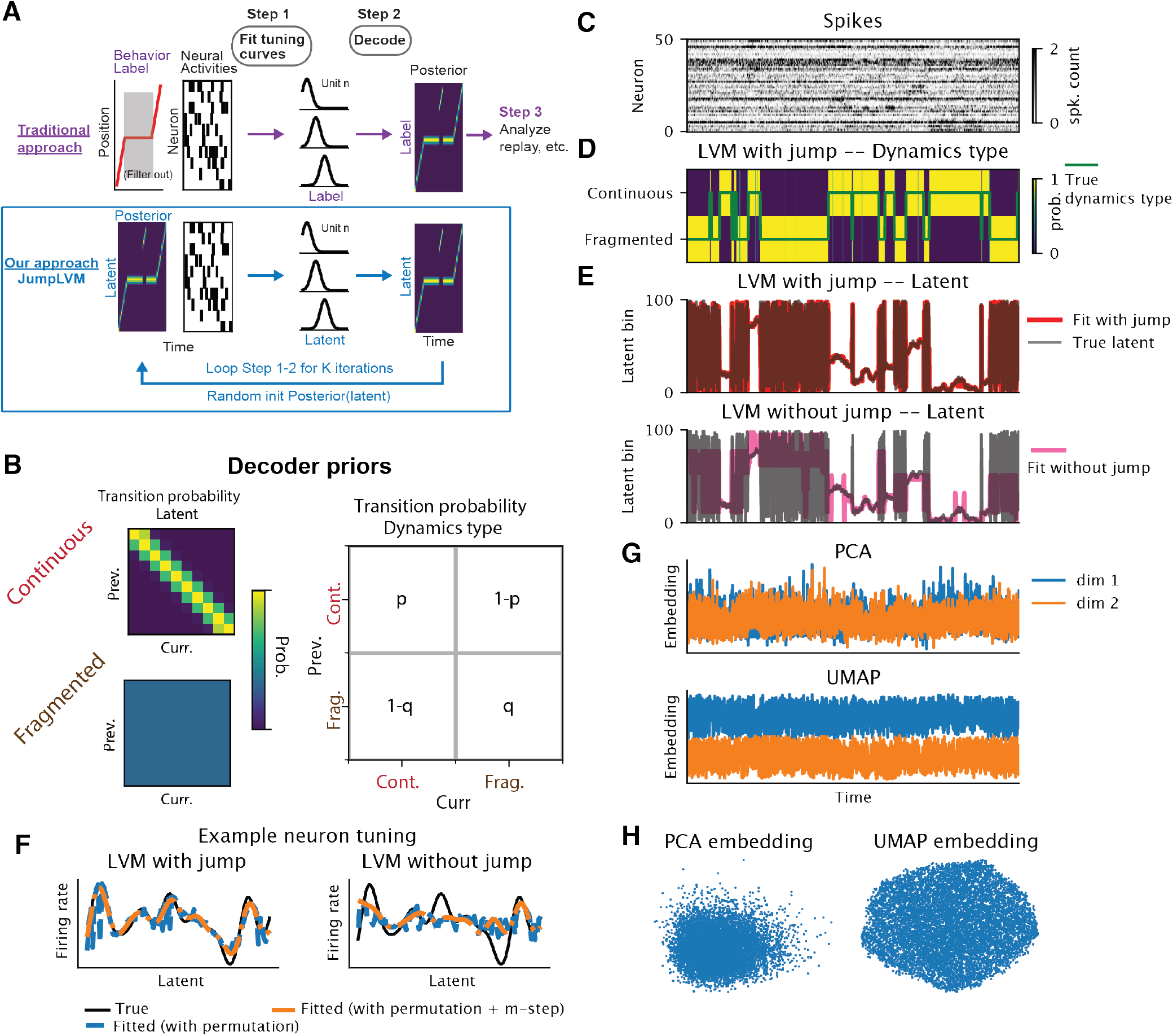
Iterative tuning-curve estimation and Bayesian decoding yield a flexible latent-variable model. **(A)** Flowcharts of traditional and JumpLVM approaches. Traditional approach (purple arrows) starts with (filtered) behavior label and neural activities, fits tuning curves, decodes and analyzes the decoded results. Our JumpLVM approach (blue arrows) starts with random (but can be label-based) initialization (init) of the latent posterior and neural activities, and iteratively fits tuning curves and decodes and, crucially, uses the result of decoding as the input for the next iteration of tuning curve fitting. **(B)** Schematics of the hierarchical priors for the decoder. Each dynamics type determines the transition probability of the latent bins. The transition probability of the dynamics types is specified by the hyperparameters. **(C)** Spike raster generated by the JumpLVM. No clear structure is recognizable. **(D)** Dynamics type, true (green line) and fitted (heatmap, more yellow color corresponds to higher probability). The inferred dynamics closely match the true values. **(E)** Latent, true (grey line) and fitted (red line, hard to see because it matches the grey line). Top: JumpLVM, Bottom: LVM without jump (and only has continuous dynamics) in the transition prior. Latent bins were permuted to best align with the ground truth. **(F)** True and fitted tuning curves for one example neuron. Left: JumpLVM recovers the tuning. Right: LVM without jump learns distorted tuning. Permuting latent bins produced the jagged tuning curves (blue), and one additional fixed-posterior M-step smoothed them (orange). **(G)** Popular dimensionality reduction methods applied to the simulated data. Both are noisy and hide the dynamics types. Top: PCA. Bottom: UMAP (compare with D) **(H)** PCA (left) and UMAP (right) embeddings in 2D. Each dot is one time bin. No structure is visible.

There are several ways to implement these modeling principles. Here, we fit neural tuning curves using generalized linear models (GLMs) over a set of pre-defined nonlinear smooth basis functions (see e.g. Pillow et al., 2008; Hardcastle et al., 2017; Balzani et al., 2020). Smoothness is essential to constrain our model, and to impose “space-like” structure on the latent space (i.e., nearby latents correspond to similar population vectors) without enforcing a one-to-one correspondence with the animal’s physical position (or any other label). For each neuron, indexed by *n*, and each time point, indexed by *t*, the GLM step yields a predicted firing rate given by *f*_*n*_(*x*_*t*_). The nonlinear tuning curve function *f*_*n*_(·) is modeled as a weighted sum of the basis smooth functions, with the weights to be optimized, and *x*_*t*_ is the input variable to the GLM. Traditionally, *x*_*t*_ is a human-supervised label (such as position or head direction). Here, we initialize a random probability distribution over *x*_*t*_ on the first iteration, but on subsequent iterations set *x*_*t*_ to the output of a Bayesian decoder (which inherits tuning curves from the previous iteration). For the Bayesian decoding step, we use the algorithm described in Denovellis et al. (2021), which discretizes the latent space into bins and uses an HMM to efficiently infer the sequence of latents *x*_1_, …, *x*_*T*_ given observed neural activity.

The most unique aspect of our model, compared to other unsupervised dimensionality reduction methods, is that it allows for not only smooth transition, but also jumps in the latent space, and infer whether the latent evolves “continuously” (i.e. transitioning into adjacent latent coordinate with high probability) or “jumps” to distant locations (i.e., transitioning into all latent coordinates with uniform probability). Smoothness is commonly imposed on the latent evolution to improve statistical efficiency and human readability (Yu et al., 2008; Wu et al., 2017; George et al., 2024). However, previous reports on fragmented decoded spatial trajectories during sharp wave ripple (SPW-R) activity, as well as abrupt changes in neural activities across brain states motivate including “jumping” (in addition to continuous) dynamics in the latent space. This functionality is achieved by our choice of the hierarchical HMM decoder developed by Denovellis et al. (2021). This model was explicitly designed to infer whether the latent trajectories evolve “continuously” or “jump” to distant locations, in addition to decoding the positions (Fig. 1B, see Methods). The smoothness in our tuning functions gives meaning to the concept of smooth transition versus jumps in the latent states. We call this new method Jump Latent Variable Model (JumpLVM). In principle, the JumpLVM model could be fit over multi-dimensional latent spaces. However, the HMM model of Denovellis et al. (2021) was developed for one-dimensional environments, and the computational costs of scaling the HMM inference and the associated latent space discretization to high dimensions are nontrivial (though not inconceivable for two or three dimensions). Furthermore, we will demonstrate that one-dimensional latent trajectories lend themselves to visualization and rigorous interpretation. Therefore, we focus on the case where the latent variable, *x*_*t*_, is a one-dimensional scalar quantity in the present paper.

After fitting the model, we obtain latent tuning curves as well as posterior probabilities of the latent variable and their dynamics. The population tuning corresponding to each latent bin captures a neural co-activity pattern. These patterns are generally more apt at describing neural data that non-linearly embed a manifold than the patterns commonly extracted from linear methods like PCA (more in Discussion, Supplementary Fig. S8). The fitted JumpLVM can then be used to decode offline activities to study reactivation and relay in an unsupervised way.

To demonstrate how JumpLVM may be used as part of a discovery tool, we simulated latent trajectories, tuning curves and spikes based on the JumpLVM generative model (Fig.1C-E). In general, establishing strict identifiability for flexible unsupervised non-linear latent-variable models remains a fundamental challenge (Khemakhem et al., 2020; Zhou and Wei, 2020; George et al., 2024). Our inference can correctly learn the dynamics type, latent and tuning curves, up to permutation of the latent bins (Fig.1E-F). The purpose of the permutation after the model fitting in this simulation is to show that we correctly identify the neural co-activity patterns that exist in the ground truth data. Since our later analysis requires only correct population patterns and dynamics, not the global arrangement of the latent space, our inference is sufficient even though the global latent arrangement is not perfectly recovered (Supplementary Fig. S1A, rows 1–2).

As one may expect, removing jumps from the JumpLVM prior results in incorrect latent trajectories and tuning curves (Fig. 1E–F; Supplementary Fig. S1A, rows 2–4). When temporal smoothness of the latent is high, the model fails to follow the jumps (Fig. 1E). When temporal smoothness is low, the no-jump model becomes too fragmented when the true latent dynamics type is continuous (Supplementary Fig. S1A, rows 3–4). Common dimensionality reduction methods, such as PCA and UMAP, fail to show any distinct structure (Fig. 1G).

However, the non-identifiability of the latent space can still induce occasional false positive jumps in the inferred dynamics type (Supplementary Fig. S1B, row 2). Moreover, real data are likely to contain elements that are mismatched to the generative model. We worried that if the true manifold is higher dimensional, fitting a 1D latent variable model would force the model to jump in some part of the latent space, even where neural activities are similar (see Supplementary Fig.S2A). We thus add a “clean-up” step after fitting the JumpLVM. The idea is that “jumps” between two states with similar tuning profiles could be spurious—i.e. arise from continuous movement along a higher-dimensional latent manifold, or non-identifiability in the model fitting. After initial fitting, we therefore impose a new continuous transition kernel that allows moving between similar tuning states without jumping (Supplementary Fig. S2B–F). We found that this clean-up step indeed removes the false positive jumps when the ground truth data do not contain jumps (Supplementary Fig. S1B, row 3). In real data, it also removes certain jumps (Supplementary Fig. S3A). However, the inclusion of the “clean-up step” only introduced minor differences in our analyses of the real data—all of the results reported in this manuscript are qualitatively similar when the clean-up step is removed (see Supplementary Fig. S3B and C).

### Continuity-fragmentation axis characterizes brain state, modulated by ACh

We first applied JumpLVM to a dataset of homecage recording of hippocampal neuron activity (*∼* 300 neurons per session) and ACh fiber photometry, during spontaneous brain state changes of wake and sleep in the home cage without behavior-tracking. We found that the dynamics type of the latent can reliably distinguish sleep states. Fig.2A shows the transition from Wake to NREM sleep, where the dominant continuous dynamics in Wake state alternates with the fragmented dynamics (i.e., latent jumps for sustained periods) and eventually settles in the fragmented dynamics regime that characterizes NREM sleep. The transition from NREM to REM shows that the fragmentation returns to continuity (Fig.2B). Overall, Wake and REM are dominated by continuity and NREM by fragmented dynamics (Fig.2E, p(continuous)>0.5). Notably the switch in the dynamics type usually precedes the switch in the brain state label from standard state-scoring algorithm based on LFP and EMG Watson et al. (2016); Levenstein et al. (2019). The higher sensitivity of the model’s prediction of brain state indicates that using spikes of neuronal populations and an appropriate model can refine the detection of brain state transitions.

**Figure 2.**
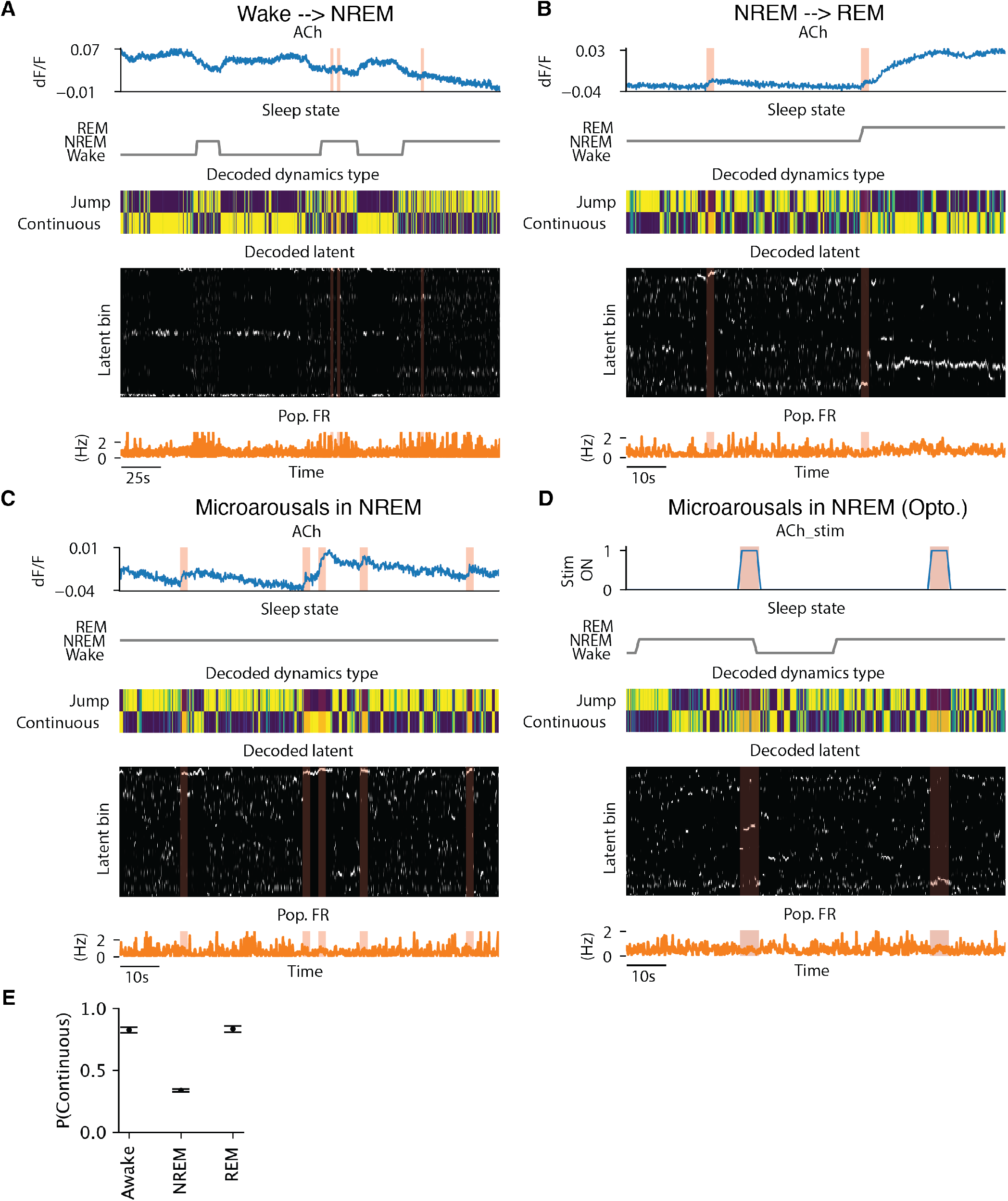
Continuity-fragmentation axis characterizes brain state, modulated by ACh. (**A-C**) Example windows of A) Wake to NREM sleep transition, B) NREM to REM sleep transition and C) microarousals in NREM. Brain state scoring is based on local field potential (LFP), and electromyography (EMG) data (Watson et al., 2016). Top to bottom: ACh fluoresence, sleep state, posterior probability of the dynamics type after the clean-up step (see *Methods*), posterior probability of the latent bin (white means higher probability), and population firing rate. Shaded boxes mark 5s windows after the detected onsets of ACh ramps that occur during NREM (see *Methods*). During wake and REM, ACh dF/F is high (1st rows), probability for the continuous dynamics type is also high (3rd rows), and the latent is more concentrated and coherent from time to time (4th rows). During NREM, the trend is mostly reversed. But bumps in ACh (shaded boxes) correlate with a shift to continuous dynamics (C). Note that the time scale between A (250 second window) and B-D (100 second window) are different. **(D)** Similar to C, except that the top panel shows optogenetic stimulation of the medial septum cholinergic neurons (The shaded boxes mark the windows of stimulation). The stimulations trigger shifts to continuous dynamics, similar to the endogenous ACh rises in C. **(E)**Average posterior probability of the continuous dynamics in different brain states. Error bars show 95% percentile bootstrap intervals obtained by resampling the means of non-overlapping 10-s chunks assigned to each brain state with replacement.

We also found that increasing levels of acetylcholine (ACh) fluorescence that characterizes NREM to WAKE/REM state changes, correlated with the continuous dynamics (Fig.2A, B). Even within NREM, when the background ACh is low and the latent evolves in a fragmented way (Fig.2C, E)), bouts of endogenous ACh rise tightly co-occurred with transient emergence of continuous dynamics (Fig.2C). To illustrate the role of Ach in the continuity regime more directly, we examined how increasing ACh release by optogenetic stimulation of the medial septal cholinertic neurons affects the fragmented dyanmics during NREM sleep. We observed immediate and consistent changes from fragmentated to continnuous dynamics (Fig.2D). Thus, JumpLVM revealed a continuity-fragmentation axis, modulated by ACh, that characterizes the population activity across sleep-wake states.

### Fragmented dynamics during wake immobility correspond to sub-threshold sharp wave ripples

We next applied JumpLVM to silicon probe recordings of the hippocampal CA1 neurons in mice (*∼* 500 simultaneous neurons) performing an alternating T-maze task (Huszár et al., 2022). The latent variable shows stereotypical sequences reflecting the trial structure (Fig. 3A, row 2). Sorting the neurons by the peaks of their latent tuning curves allowed us to visualize the spike raster in a way that confirm the structure of the latent variables (Fig. 3A, row 3). The latent variable model explains the held-out data considerably better than the supervised tuning curve model (Fig.3B). Specifically, we compared all models to a baseline model that assumed a constant firing rate (homogeneous Poisson process), which explains -4.2 (movement) and -4.8 (immobility) bits-per-spike on average across neurons. Relative to this baseline, a supervised tuning curve model explains an additional 0.76 (movement) and 0.73 (immobility) bits-per-spike on average across neurons. Depending on the hyperparameters, the latent variable models explain somewhere between 0.90 and 1.19 bits-per-spike over the baseline, representing between 14% and 55% additional bits-per-spike over the supervised tuning curve model. The jump model also achieves up to an extra 8% gain compared to the continuous-only model (Fig. 3B). When smaller time bins were used (20 ms instead of 100 ms), the performance gain over the continuous-only model was larger, approaching 30% (Supplementary Fig. S4). Downsampling the number of neurons did not hurt the test performance on average, but increased the variability (Supplementary Fig. S7A).

**Figure 3.**
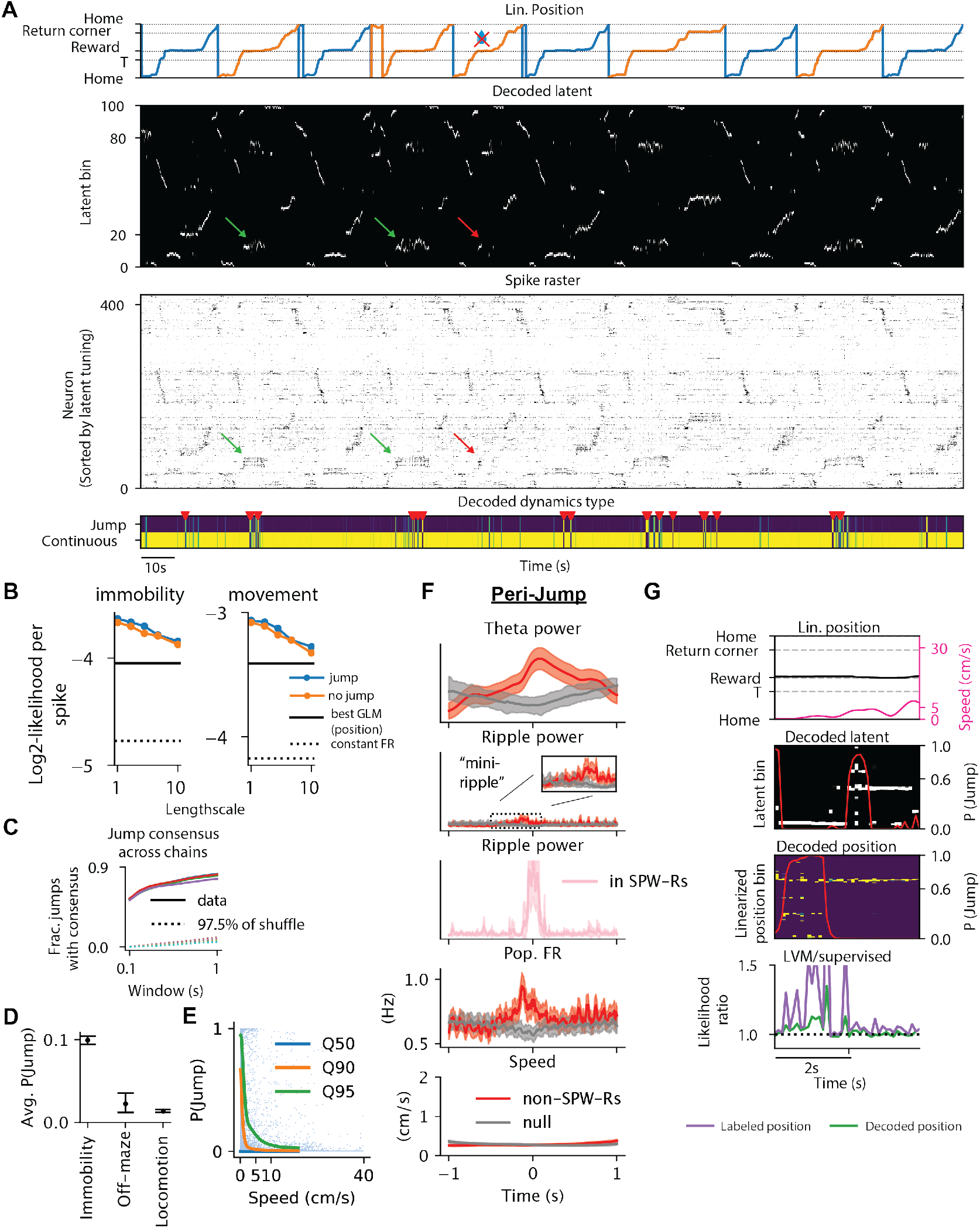
Latent jumps occur during wake immobility, accompanied by mini SPW-Rs. **(A)** JumpLVM results on multiple trials of T-maze alternation. Row 1: linearized position. Different colors indicate different turns. The no water symbol indicates an error trial. Row 2: Posterior probability of the latent in time. White indicates high probability. The green arrows point to latent when the animal was stationary at the correct reward port. The red arrow points to same latent as green, but when the animal was at the incorrect reward port. Notice the very brief activation of the previous latent during the error trial, followed by different latent activation. Row 3: Raster plot of population spike counts. Neurons are sorted according to the peak of their latent tuning curves. Arrows as before, but now pointing at the corresponding subset of neurons. Row 4: posterior probability of the dynamics type in time, after the clean-up step (see *Methods*). Yellow indicates high probability. Red triangles mark the the consensus jumps. **(B)** JumpLVM out-performs both the spatial baseline and the model without jump at different lengthscales, during both movement and immobility. Larger lengthscale corresponds to higher smoothness in the tuning curves (in the unit of latent bin). Y axis: log_2_ -likelihood-per-spike (i.e. log likelihood normalized by total spike counts) for different models. For the JumpLVM (blue) and LVM without jump (orange), we display the normalized log marginal likelihood on held out data, split by speed threshold, as a function of the tuning curve lengthscale. The black dotted line is the inhomogeneous Poisson (i.e. constant firing rate) baseline model. The black solid line is the best generalized linear model (GLM) with a position regressor (selected over different regularization strengths and numbers of basis functions)). **(C)** Fraction of jumps with consensus as a function of window size, demonstrating the robustness of the jump detection. Putative jump times are defined by *P* (*Jump*) *>* threshold. The threshold varies from 0.3 to 0.7; thus the choice is not critical (different colors overlay; threshold is fixed to be 0.6 in all other analyses). Consensus for each jump is defined as four out of the five random initializations having putative jumps within ± window relative to that jump (fixed as 0.5*s* in all other of analyses). The dotted lines are the 97.5% of the shuffled distribution of fraction of consensus jumps, obtained from circular shuffles of the *P* (*Jump*) time series independently for each random initialization. **(D)** Average jump probability during each behavior type. “Off-maze” refers to when animal’s position tracking deviates from the maze, likely reflecting head-scan and rearing behaviors (see Fig. 4B). Error bars show 95% percentile bootstrap intervals obtained by resampling the time points with replacement. **(E)** 2D Histogram of jump probability (Y axis) and running speed of the animal (X axis). The colored lines mark different percentiles of jump probability at each speed level. High jump probability tends to occur at low speed. **(F)** Jump event-triggered averages correlate with sub-threshold SPW-Rs. The red curves are from jump events that occur outside of detected SPW-Rs and during immobility, while the pink curve (third row) is from events that occur within detected sharp-wave ripples. The shades are 95-CIs. The grey curves are derived from time bins during immobility without jumps and SPW-Rs. Pop. FR, average population firing rate. **(G)** Example window where JumpLVM and supervised spatial decoding show opposite dynamics, with JumpLVM showing superior performance explaining spikes. Row 1: linearized position and speed. Row 2: posterior probability of the latent. Jump probability in red line. Row 3: posterior probability of the decoded position from supervised spatial decoding. Jump probability in red line. Row 4: likelihood ratio between the JumpLVM and supervised models. The likelihood for each model is computed from the expected firing rates and the Poisson observation. For the labeled position (purple), expected firing rates are computed by evaluating the tuning curves at the given position label. For the decoded position (green) and the JumpLVM, expected firing is the posterior weighted average of the tuning curves evaluated at each position / latent bin.

The benefit of JumpLVM is not merely quantitative. It reveals that, during an error trial, both the latent variable and the spiking activities are distinct from the preceding correct trials when the animal visited the same reward port (Fig. 3A, green and red arrows, Supplementary Fig. S5). In contrast, supervised spatial decoding decodes the same physical position (as it is supposed to do), failing to reveal the different neural activity patterns during the error trial, while having a low likelihood in explaining the spikes (Supplementary Fig. S5C-D).

We next sought to validate that the jumps during wake behavior learned by the model are reproducible and scientifically interpretable. The clean-up step (Methods) is important here to reveal that many seeming gaps in the latent are in fact continuous based on the tuning similarity (Fig. 3A, row 2 and 4; Supplementary Fig. S3A, row 3 and 4). We observe that when we fit the JumpLVM model multiple times from different initializations and performed the clean-up step, the different models “agree” on a large fraction of the jump times. For example, when we fit five models on the alternating T-maze dataset, 73 percent of jumps were found across at least 4 of the models within 500 ms tolerance. Regardless of how we set the threshold for what is considered a jump and the window size for defining consensus, we found that the amount of consensus across models was significantly higher than shuffle, and was as high as 55% within ± 0.1 second (Fig.3).

Thus we studied these “consensus jumps” further to if these events defined from spikes are interpretable in terms of behavior and LFP. Consistent with our previous result that jumps occur more during NREM sleep, in this spatial navigation task we also saw higher jump probabilities during immobility compared to other behavior types (Fig. 3D-E).

To examine physiological signatures associated with jumps while minimizing movement-related confounds, we included only jumps for which the animal’s speed was below 1 cm*/*s during the latent jump. While it is reasonable to suspect SPW-Rs to underlie the jumps, we actually found the opposite. 92.8% (543 jumps during slow speed time bins and outside of SPW-Rs / 585 jumps during slow speed time bins) of the consensus jumps during immobility did not occur during SPW-Rs. However, within a small time window before those jumps, there is a small but significant increase in the ripple band (100-250Hz) power and population firing rates (Fig.3F). The power increase is minisculous compared to the jumps that occurred within SPW-Rs (Fig.3F, row 2 vs 3). We thus call them “mini-ripples”. This finding highlights the potential of the model for identifying physiologically meaningful events from spiking data alone.

Given that fragmented dynamics were first used in supervised spatial decoding (Denovellis et al., 2021), it is reasonable to question whether the supervised spatial decoding is sufficient to characterize continuous vs fragmented dynamics when spatial labels are available. Fig 3G shows an example where the JumpLVM and supervised state-space decoder predict fragmented dynamics at opposite times, with the JumpLVM predicted firing rate explaining the spikes better than the firing rate predicted from both the position label or the decoded position (the latter incorporated information from the population and is thus harder to beat).

### Behavioral state classifies latent variable

The JumpLVM model identifies a 1D latent manifold discretized into 100 equally spaced bins. We next sought to interpret the latent variable both in terms of its manifestation on the maze and its reactivation during sleep before (PRE) and after (POST) the task (Fig. 4A-C). At each moment in time, the model maps the neural activity onto a probability distribution over these “latent bins”, according to their consistency with the data. We then selected the maximum a posteriori (MAP) latent bins at each time. Projecting these latents onto the physical environment revealed a spatial organization of the latents, a population analogue of “place fields” (Fig.4C, top rows, Maboudi et al. 2018). This visualization makes it apparent that the population spatial representation drifts within a session (Fig.4C, latent 63, blue to red expressed at different physical locations, Yang et al. 2024; Zheng et al. 2024). It also highlights differences in the precision of the spatial representation on different parts of the maze (O’Keefe and Speakman, 1987; Hollup et al., 2001; Burke et al., 2011; Sato et al., 2020). For instance, latent 12 shows a narrow field, while latent 99 has a much wider field (Fig.4C).

**Figure 4.**
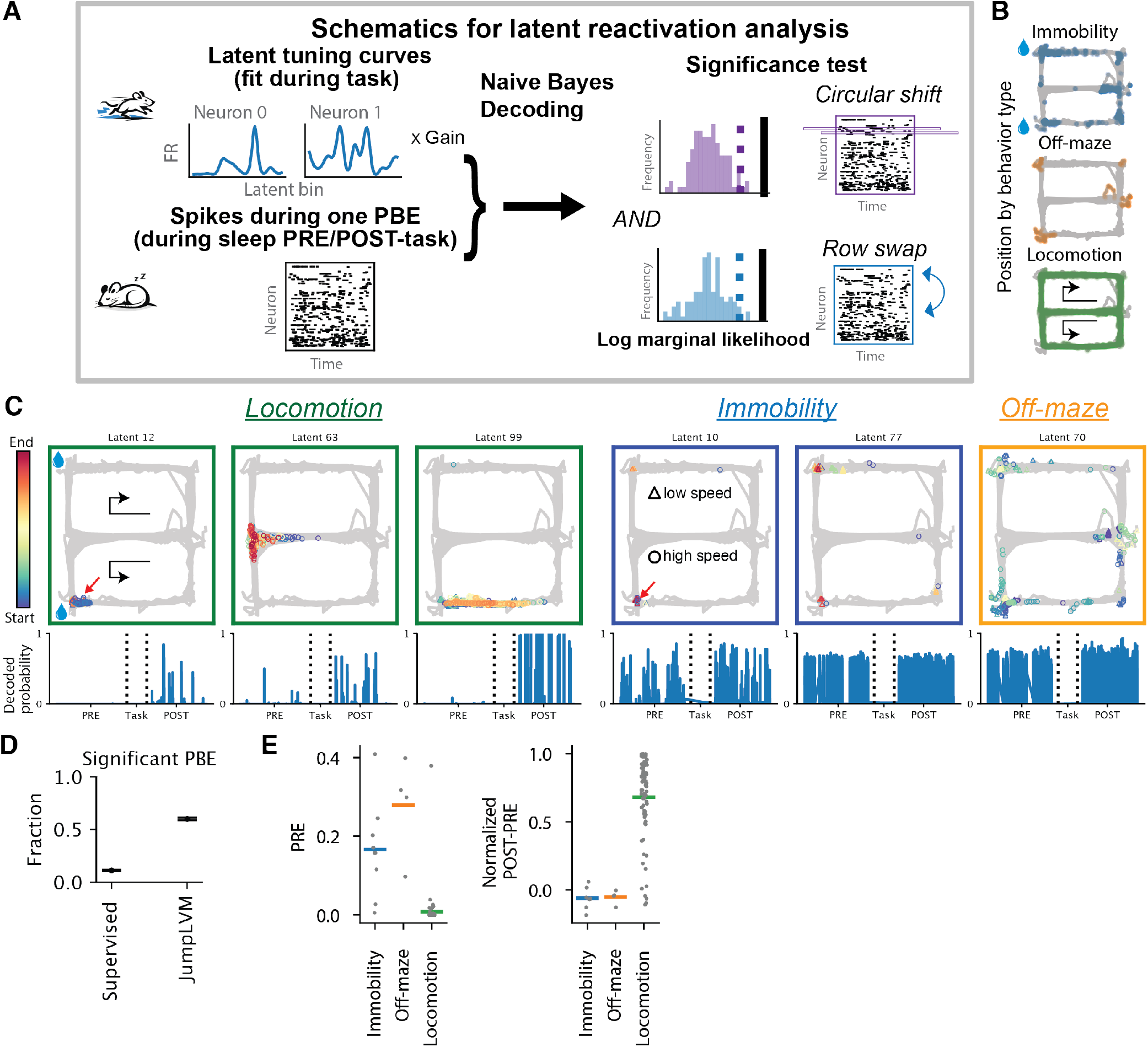
Latents show spatial tuning and uncover differential reactivation based on behavioral state. **(A)** Schematics of the latent reactivation analysis. **(B)** Illustration of different behavior types. The background grey lines are the mouse’s trajectory throughout the whole session, delineating the maze. The colored dots mark the mouse’s positions during each behavior type. **(C)** Latent place fields and reactivation in NREM sleep. Top row: each dot is the position of the animal on the maze when the given latent achieves the maximal posterior, colored by elapsed time in session, and shapes determined by speed (5cm/s speed threshold). The color of the surrounding box indicates the the behavior type of each latent. The bottom of each column is the posterior probability during significant population burst events (PBE, re-binned at 20ms) during NREM sleep throughout the recording session. Note increased incidence of locomotion-related (Beh) latent during post-task (POST) sleep versus pre-task sleep (PRE). X axis is time. Vertical dotted lines indicate the beginning and end of the maze navigation. **(D)** Fraction of PBEs that are significant reactivations. Error bars are 95% percentile bootstrap intervals obtained by resampling the PBEs with replacement. **(E)** Immobility and off-maze latents have higher pre-activation and locomotion latents have higher reactivation. Left: quantification of the pre-activation: average posterior probability of each latent type during PRE-task sleep (max within each significant PBE, mean across them, same for right); right: reactivation: measured as (POST-PRE)/(POST+PRE) average posterior for each latent type (immobility and off-maze, locomotion). Each dot is from a latent bin. Horizontal bars mark the mean across latents. Note that only locomotion-related latent bins have low activation before the task (PRE) and increase after the task (POST).

However, “coding of space” is not enough to explain all the latents. To our surprise, some latents occur at multiple parts of the maze, but mostly when the animal’s trajectory deviates from the maze, potentially during head-scanning and rearing but not ambulating behaviors (which we call “off-maze” (Fig. 4B-C). Furthermore, while the latent 12 and latent 10 (Fig. 4C, top) have similar place fields, the former mostly occurs during locomotion in the maze (circle markers), while the latter mostly occurs during immobility and drinking (triangle markers). These observations prompted us to classify the latents into “locomotion”, “immobility” or “off-maze” (rearing/head-scan) (see Method) based on the behavioral types of the animal when each latent mostly occurs. Interestingly, some of the immobility latents occur mostly at one port, with sparse activation at the other port (Fig. 4C, top, latent 10), other immobility latents are similarly active at both reward ports (Fig. 4C, top, latent 77), potentially reflecting a general “reward-related brain state” (Gauthier and Tank, 2018).

### Behavioral state modulates latent reactivation

Given the behavior-modulation on the hippocampal population activities, a natural question is how these different categories of latents are reactivated during NREM sleep. Using the learned tuning curves from the JumpLVM as the templates (multiplied by a gain, see Methods), we performed Naive Bayes decoding on the binned spike counts during population burst events (PBE, see Methods) and used the log marginal likelihood as a measure of how well the tempalte explains the data. We performed neuron id and circular shuffle on the PBE spiking matrices (van der Meer et al., 2020; Yang et al., 2024) and the Naive Bayes decoding to get a null distribution of the log marginal likelihood. A PBE is considered a significant reactivation if its log marginal likelihood exceeds the 95-percentiles of both shuffles (Fig. 4A). We performed the same procedure using the supervised spatial tuning as the template. JumpLVM discovered a 60% of significant events, whereas the supervised spatial decoding only 11% (Fig. 4D). The 6-fold increase strongly suggests that the co-activity patterns induced by spatial labels are misaligned with the internally generated co-activity patterns during sleep. These endogenous patterns are much better captured by JumpLVM since it extracts templates based on the non-linear manifold structure of the neural activities. Downsampling the neurons reduced the fraction of significant PBEs, suggesting the importance of the number of simultaneously recorded neurons in identifying reactivation (Supplementary Fig. S7B, Tingley and Peyrache, 2020; Takigawa et al., 2024; Yang et al., 2024).

Strikingly, the different classes of latents have different patterns of reactivation (Fig.4C, bottom), supporting the biologically meaningful nature this latent classification. The locomotion latents show classic reactivation, with low occurence in PRE-task sleep and high incidence during POST-task sleep (Fig. 4C, bottom and E). In contrast, the immobility and off-maze latents occur at higher frequency but their incidence does not change between PRE-task and POST-task sleep (Fig. 4C, bottom and E). This pattern is robust across a wide range of hyperparameters (Supplementary Fig. S6).

### Analysis of non-spatial replay via decoding of dynamics type

The preceding reactivation analysis only answered the question of whether the pattern of neural activities within one time bin repeat during offline time bins. Do the temporal dynamics of the reactivation also follow the same sequential structure as during the Task epoch, especially for those non-locomotion behaviors whose replay have not been studied (e.g. off-maze)? We answer this question by equipping JumpLVM with a new set of dynamics type, i.e. latent transition probabilities (Fig. 5A). These dynamics types (immobility, off-maze and locomotion) were motivated by the behavior types that were found to be important categories from the latent analysis in the previous section. After fitting the JumpLVM using the prior latent transition, consisting of the continuous and fragmented dynamics types, We computed the empirical latent transition probability separately during the three behavior types: immobility, off-maze and locomotion. The empirical latent transitions largely match the continuous dynamics in the prior, but also show crucial deviations, particularly for the off-maze (Fig. 5A). We replaced the continuous dynamics in the prior with these behaviorally derived transitions. Together with the same fragmented dynamics, they form our new model of latent transition, which we used to decode the spike count matrices during significant PBEs. We then applied a threshold on the posterior probability of the dynamics types averaged within each PBE to classify the PBE into replays of different behavior types (or unclassified) (Fig. 5A).

**Figure 5.**
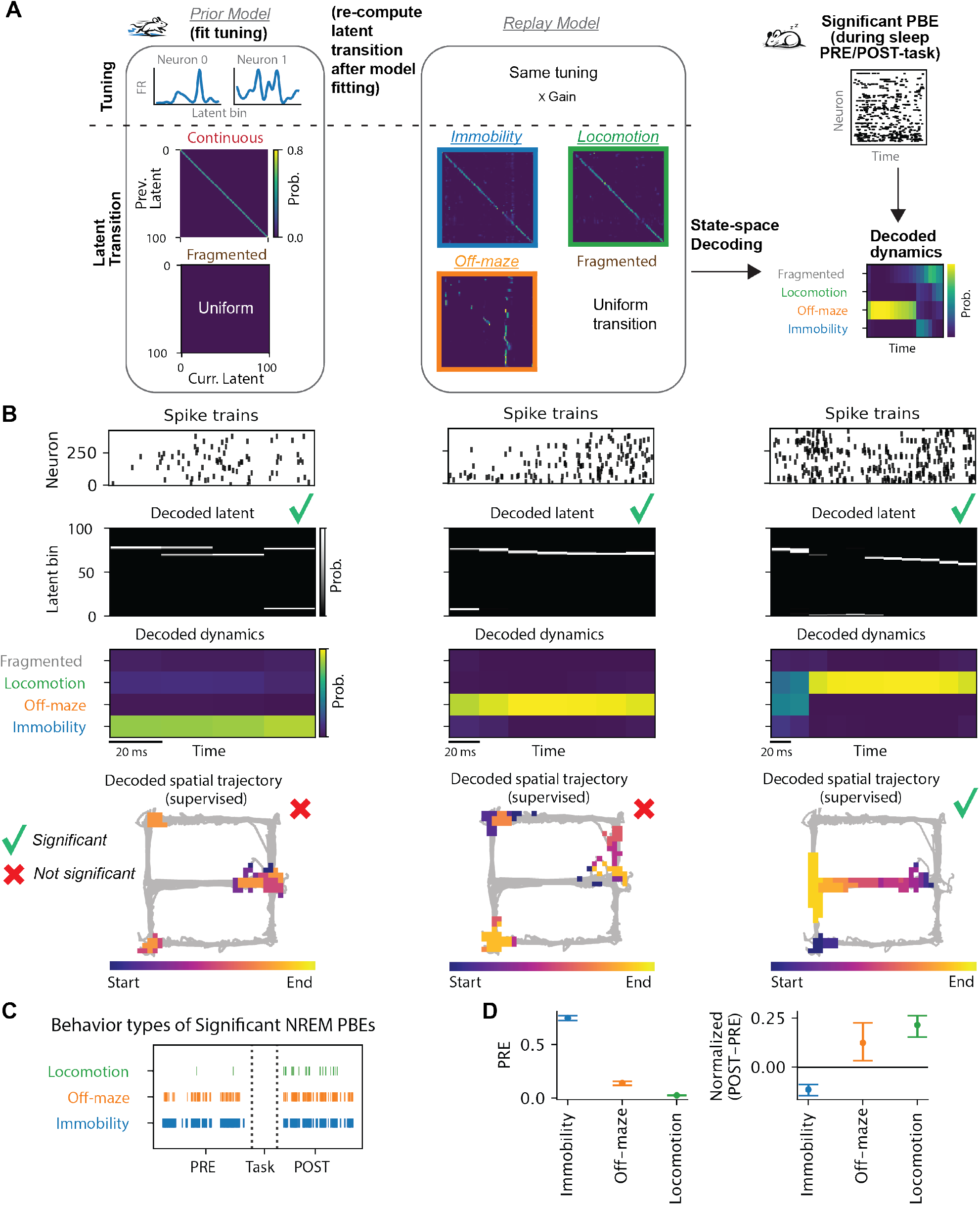
Replay of different behavior types. **(A)** Schematics of the JumpLVM replay analysis. After fitting the model to the Task epoch, we recomputed latent transition probability using the posterior probability of latents within each behavior type of interest. These probabilities replace the continuous prior transition, and constitute the dynamics types together with the fragmented dynamics for the replay model. State-space decoding was performed using this replay model to determine the dynamics type of each moment of the PBEs. Only significant PBEs (Fig. 4) A) are used for the replay analysis. **(B)** Example replays. Top to bottom: spike rasters; posterior probability of the latent bins; posterior probability of the dynamic types; high posterior bins (above 90-percentile at each time bin) from supervised spatial decoding, colored by progression within the event. Columns from left to right are examples for immobility, off-maze and locomotion replays (Note the events have unequal durations). The green check marks on the second row shows they are all significant PBEs under the JumpLVM derived template, while the red crosses on the bottom row of the first two columns indicate they are not significant under a supervised spatial template. The supervised decoding (third row) qualitatively supports the replay classification based on the JumpLVM dynamics, but lacks precision and statistical power. **(C)** Classified replay occurrences (mean probability for one dynamics type within PBE higher than 0.8). X axis is time. Dotted vertical lines mark the start and end of the Task epoch. **(D)** Left: probability of each dynamics type averaged within PBE, median across significant PBEs during the PRE-task NREM sleep. Right: the same aggregation process as the left to get POST and PRE, but plotting the measure of replay: (POST-PRE)/(POST+PRE). Error bars from bootstrapping.

Fig. 5B shows example replays classified by the JumpLVM. The spike rasters (Fig. 5B, top row) highlight the difficulty of seeing replay by naked eyes without the help of statistical modeling, when number of neurons are large. While the decoded latents seem to have large jumps (Fig. 5B, second row), they are largely consistent with one of the three behaviorally derived dynamics types and not with the fragmented dynamics (Fig. 5B, third row). For the event classified as locomotion (Fig. 5B, third column), the supervised spatial decoding (Denovellis et al., 2021) shows a long coherent trajectory. Crucially, for the immobility (Fig. 5B, first column) and off-maze replays (Fig. 5B, second column), the events did not pass the shuffle test using the supervised spatial template, even though the spatial decoding suggests a replay content consistent with the behavior type classified by JumpLVM. For instance, the immobility replay indeed had stationary (albeit multimodal) decoded positions (Fig. 5B, first column, fourth row). The off-maze replay also had much posterior mass off the maze (Fig. 5B, second column, fourth row). The fact that these events fail the significance test (under the spatial template) and the multimodality of the decoded position might disqualify them as replays based on common space-centric criteria. However, here we show they are bona fide reinstatement of neural activity patterns following certain temporal dynamics during particular wake behaviors. A final interesting detail that can be gleaned from this type of replay analysis is that there can be a transition of behavior types within an event. The locomotion event had high probability for off-maze dynamics at the beginning of the event, which is also hinted by the spatial decoding, but could easily be dismissed as noisy multimodal decoding at the start of the trajectory (Fig. 5B, third and fourth rows).

The distinct patterns of reactivation from the previous section is largely recapped by the replay analysis. Immobility and off-maze events are pre-played frequently, while the locomotion replays occur mostly during POST-task sleep (Fig. 5C-D). These effects are robust under a wide range of downsampling of the neurons (Supplementary Fig. S7C). The pre-existence of immobility replay and the prominence of locomotion replay are also robust under a wide range of hyperparameters (Supplementary Fig. S6). When the tuning lengthscale is low (i.e. wiggly tuning curves), the off-maze replay becomes not prominent during PRE-task (Supplementary Fig. S6B, column 2). However, small tuning lengthscales would over-split the latent, making the transition estimation unreliable (Supplementary Fig. S6C, column 2), justifying our choice of a smooth lengthscale.

### JumpLVM identifies theta flickering in MEC without position labels

Finally, we demonstrate how the JumpLVM can be used for exploratory analysis of high-dimensional datasets. Vollan et al. (2025) used a variant of the Poisson GPLVM (Wu et al., 2017) on neuropixel recordings from the medial entorhinal cortex (MEC, *∼* 800 units) and showed that the population spatial representation swept off the linear track into the untraversible space and alternated sides on a theta timescale. To identify such theta sweeps typically requires initializing the latent with spatial labels, and especially, in 2D, which is not an intuitive starting point for 1D track. We show that even without any additional knowledge (i.e. no spatial label and no 2D latent), the JumpLVM model can detect theta flickering. Applying the JumpLVM to the spikes binned at theta time scale (100ms, as opposed to 10ms in Vollan et al.,2025), we see fragmented dynamics during traversal of the linear track, where the latents progress in two lines, jumping back and forth at each time bin,i.e. theta time scale (Fig. 6). This demonstrates that the population firing pattern is similar to every other theta cycle (cycle skipping, Kay et al.,2020). To reach this conclusion, our procedure required twenty seconds of computation, in contrast to the original procedure that requires hours.

**Figure 6.**
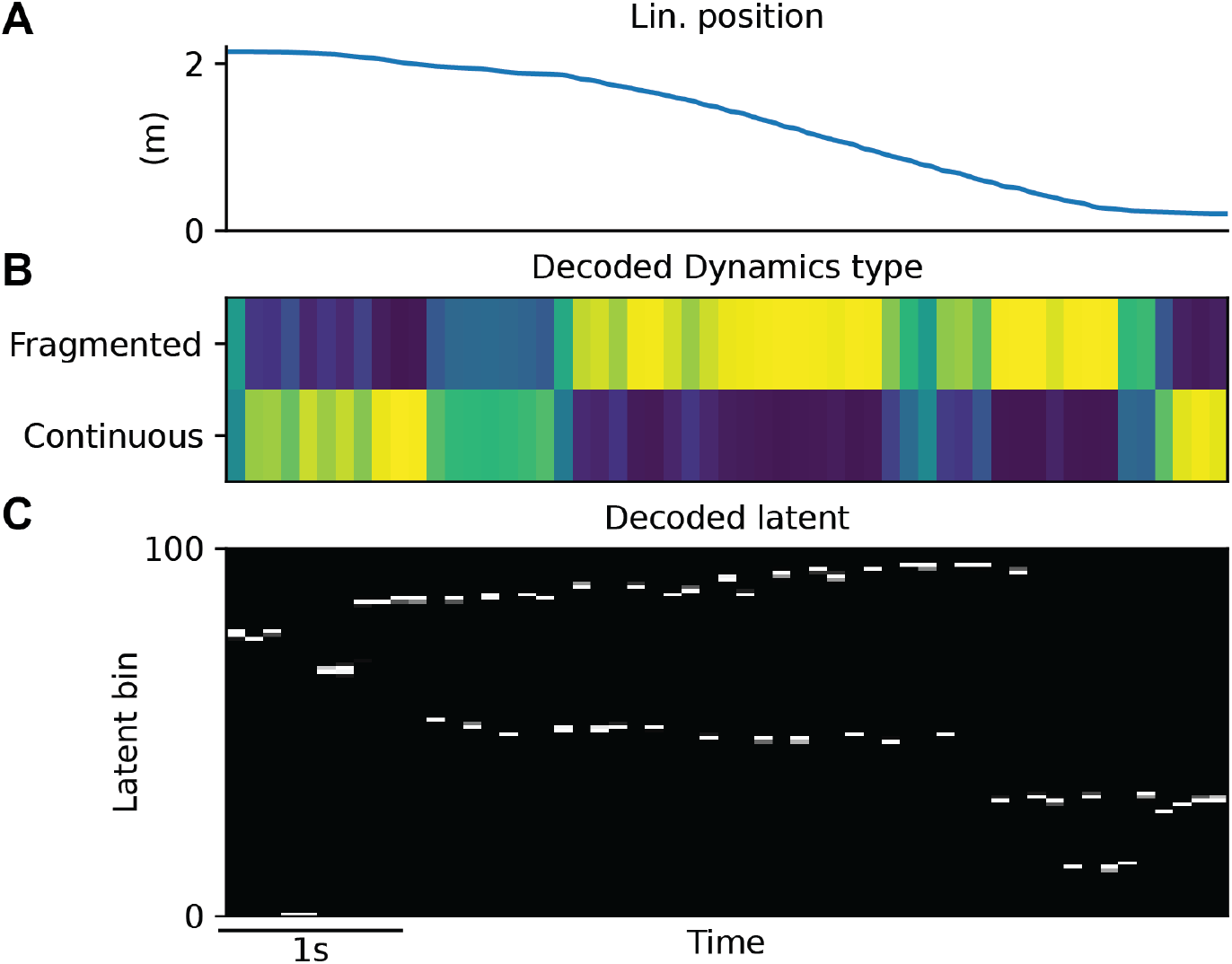
JumpLVM readily identifies theta-flickering in MEC. **(A)** Position on a linear track of one example trial. **(B)** Decoded dynamics type is dominated by fragmented but periodic dynamics during the run. More yellow in color means higher posterior (same as in C). **(C)** Decoded latent at theta time scale (100ms bin) exhibits flickering during the run.

## Discussion

We developed JumpLVM, a flexible latent variable model for discovering structure in neural population activity without requiring behavioral labels as the starting point. Our central finding is that hippocampal population activity is organized by behavioral and brain-state variables that are only partly captured by spatial labels. In a spatial navigation task, JumpLVM identified latent activity patterns associated with locomotion, immobility, and off-maze behaviors, and revealed that these patterns are reactivated and replayed differently during NREM sleep. Furthermore, by allowing the latent state to evolve either continuously or through jumps, the model uncovered a continuity–fragmentation axis that characterizes hippocampal dynamics across wake and sleep.

JumpLVM has general applicability in any settings where neural activity may reflect a mixture of smooth evolution and abrupt state changes. JumpLVM could be useful for identifying internal states not obvious from behavior, parsing how action policies, habits, or internal task states are represented and replayed.

### Jump in latent variable models

Most latent variable models assume neural activity changes smoothly over time, such as GPFA, GPLVM, LFADS and SIMPL (Yu et al., 2008; Wu et al., 2017; Jensen et al., 2020; Pandarinath et al., 2018; George et al., 2024), to name a few. In those models, the latent state is encouraged to move gradually, so nearby timepoints are represented by nearby latent values. That works well if the underlying neural process is genuinely continuous, but it can blur abrupt transitions. One potential worry that prevents the detection of abrupt changes in latent variable models is that the lack of the smoothness prior might subject models to overfitting. We found a way to keep the smoothness prior most of the time, while allowing for occasional jumps, thus avoiding the potential overfitting. We verified that jumps are not merely model artifacts, but reflect genuine large changes in spiking activity, supported by additional physiological markers. We found that sustained jumps co-occur with “mini” SPW-Rs. Including fragmented dynamics also provides an interpretable axis for characterizing population activity across wake-sleep states, as well as the effect of cholinergic modulation on hippocampal circuits. These observations illustrate and justify the need of including discontinuities in our assumptions about the dynamics of neural activities. This opens up the opportunity to study abrupt changes in other systems, for instance, during decision making, movement initiation or motor action switching.

Identifying moments of continuity and discontinuity can provide physiologically meaningful windows for further examination. For instance, our JumpLVM identified epochs of continuous dynamics amidst the background of fragmented dynamics during NREM, which could serve as a proxy for neuromodulator increases (arousals). Furthermore, while our model was not designed for classifying sleep states, we found clear signatures of sleep state transitions based on the latent dynamics, which could be exploited to improve the accuracy of brain state scoring in the future.

### JumpLVM as non-linear assembly detection

Common non-linear dimensionality reduction methods, like Isomap, t-SNE and UMAP (Tenenbaum et al., 2000; Maaten and Hinton, 2008; McInnes et al., 2018) excel at visualizing the global structure of the data, e.g. rings in manifolds. JumpLVM is not another method for visualizing the low-dimensional embedding. It is better treated as a non-linear assembly method. The population tuning with respect to each discretized latent bin can be regarded as a co-firing assembly. The decoded probability of a latent bin can be viewed as activation strength. When projecting the latent onto the maze, we discovered latent bins that are active during immobility and off-maze behaviors. Notice that the discovery of off-maze latent would be not intuitive using the common way of reducing the population activity to 3D scatter plots colored by behavior labels like position (Yang et al., 2024; Sun et al., 2025). Instead, our approach projects the discretized latent onto animal’s trajectories on the maze. The fact that some latents occur as the animal’s position appears off the maze and not during locomotion naturally suggests the distinctness of these events.

### Analysis of non-spatial replay

Most hippocampal replay analysis assumes a spatial template (Diba and Buzsáki, 2007; Karlsson and Frank, 2009; Foster and Wilson, 2006; Tingley and Peyrache, 2020; Denovellis et al., 2021), and report a small fraction of offline PBEs containing significant replay events. We showed that this could come from the fact that spatial templates are not the optimal axis for interpreting the hippocampal off-line activities. In fact, we found a 6-fold increase in the number of significant off-line reactivation events when we replaced the spatial template with the unsupervised template learned from JumpLVM. This suggests reactivation and replay are more general and prominent feature of hippocampal function than previously appreciated. In addition, JumpLVM highlights the categorical difference between the contents that are preplayed (Dragoi and Tonegawa, 2011; Farooq and Dragoi,2019) (immobility and off-maze behaviors) with those that are mostly replayed (locomotion). The perpetual activation of the immobility and off-maze latents during sleep echos the predominance of the stationary spatial replay during wake immobility in Denovellis et al. (2021). Neurons involved in these latents potentially form a backbone (Grosmark and Buzsáki, 2016), whose function is up for future investigations.

### Comparison with alternative unsupervised methods for reactivation and replay analysis

The non-linearity of our method also separates it from linear assembly detection methods based on PCA and independent component analysis (ICA) (Lopes-dos Santos et al., 2013; Van de Ven et al., 2016; Huszár et al., 2022; Liu et al., 2023; Zutshi et al., 2025; Peyrache et al., 2009, 2010; Gulati et al., 2014; Ramanathan et al., 2015; Gulati et al., 2017; Kim et al., 2019; Tingley and Peyrache, 2020). The assumption of the assembly detection is that linear modes of the correlation matrix reflects co-activating cell assemblies. While this assumption is reasonable in brain regions where low dimensional subspaces capture a large variance (Churchland et al., 2012), it becomes questionable when the neural activities embed a curved manifold in a non-linear fashion (De and Chaudhuri, 2023), e.g. in the hippocampus (Rubin et al., 2019). However, linear modes reflecting global correlational structure can be distinct from the nonlinear manifold structure (for a toy example demonstrating the issue, see Supplementary Fig. S8).

Furthermore, the inclusion of temporal dynamics also separates our approach from these other reactivation analysis of non-spatial variables, which treat time as independent. Hidden Markov Model (Rabiner, 2002, HMM) is similar in terms of using transition probability, which was fitted to PBEs for replay analysis by Maboudi et al. (2018). Our method can be thought of as an HMM with fixed transition matrix and smoothed observation. The advantage of JumpLVM is that the fixed transition guide the model to learn good solutions more easily, while HMM is prone to local minima (Rabiner, 2002; George et al., 2021; Li and Camera, 2025), especially when the number of latent bins (or “state” in the HMM terminology) is large. The learned latents immediately offer a visualization of sequences and jumps without additional post-hoc sorting. More importantly, instead of asking whether the offline activity reply one transition graph as in HMM (“replay or not”), our approach classifies the PBEs into interpretable types (“locomotion or immobility or off-maze or unstructured).

### A complementary role for unsupervised methods

With the increase in the number of simultaneously recorded units, there has been a rapid growth in unsupervised statistical methods for discovering structure in neural population activity. Much of this work has focused on settings where there is a strong prior understanding of what the circuit encodes, and has evaluated models by asking whether latent variables recover known task variables. For example, latent manifolds in hippocampal recordings often align with physical space, and latent trajectories in motor and premotor areas with reach kinematics. Newer models are frequently benchmarked by improved spike prediction or decoding performance relative to earlier approaches. Demonstrating that these methods extract sensible structure consistent with prior studies, and that they explain the data well, is an essential step in validating the approach.

At the same time, the distinctive strength of unsupervised approaches is their potential to reveal structure when we do not already know what a system encodes, or to uncover additional variables beyond the experimenter’s label set. In this spirit, the present work emphasizes what can be learned when we reduce our reliance on prior behavioral labels, or when such labels are not available. Even in a task with clear labels and well-characterized tuning (spatial navigation and CA1 pyramidal neurons), the latent variables inferred by JumpLVM provide a richer description of the underlying population activity. For example, CA1 activity patterns at the same position differ depend on other task variables (e.g., correct vs. incorrect) as well as behavioral state changes (e.g., running vs. off-maze head scanning). These observations suggest an expanded “bank of labels” for subsequent supervised analyses. Furthermore, classifying latent dynamical regimes offered us to identify “mini” sharp-wave–ripple–like events underlying fragmented dynamics, and yielded an interpretable continuity– fragmentation axis that characterized population activity across wake and sleep, modulated by acetylcholine (ACh). Together, these examples illustrate how JumpLVM can facilitate biological discovery.

### Limitations of the model

The model does not explicitly account for the distributional shift in firing statistics that can occur during SPW-Rs. We instead resort to fitting the model with a coarser bin (100ms) and decoding SPW-R content with a finer bin (20ms), together with a gain on that tuning that is selected post-hoc. A future direction is to fit all the data simultaneously, without splitting data into “template” and “match.” To do so the model has to infer a distributional shift automatically and extracting latents evolving at multi-timescales simultaneously. While using the model as a non-linear assembly detection is fruitful in the currently analysis, it would be useful to be able to reliably recover the topology among the latents.

## Acknowledgment

We thank everyone from the Williams lab and the Buzsaki lab and Jonathan Pillow for their helpful feedbacks and discussions. This work is supported by NIH/NIMH RF1MH133778, MH13916 and U19NS107616. We also thank the McKnight Foundation.

## Methods

### Generative model

The jump latent variable model assumes that the neural activities *Y* (time by neuron) are generated by latent variables *X* (time by num. latent bin) given the tuning curves *F*_*n*_ per neuron and the noise model (for spikes, we use the Poisson distribution). To add smoothness to the latent tuning curves, we model them as Gaussian

Processes (GP) on the latent space (Williams and Rasmussen, 2006). The smoothness biases similar latent coordinates to have similar population responses, thus imbuing a sense of “closeness” to the latent space. However, naive Gaussian Processes require *O*(*T* ^3^), which is prohibitive when *T* is more than a few thousands (see Liu et al.,2020 for a review of scalable GP). We therefore use smooth basis functions *{ϕ*_*b*_(*·*) *}* _*b*_ and weights *W* to model latent tuning (similar to Pillow et al.,2008;Williams and Rasmussen,2006) (*t*: time, *n*: neuron, *b*: basis):

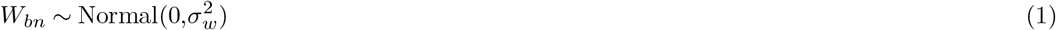

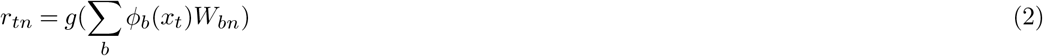

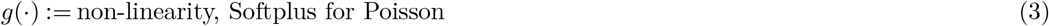

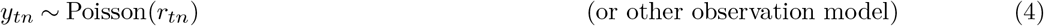

The latent variables *x*_*t*_ transition in time under either a gaussian random walk prior (with variance 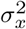) or a uniform prior, determined by the dynamics type *I*_*t*_ (‘continuous’ or ‘fragmented’/’jumping’) (In the paper we use ‘fragmented’ when the jumps occur in an extended period of time). The dynamics follow a Markov chain specified by the transition probability matrix *M* (Denovellis et al., 2021):

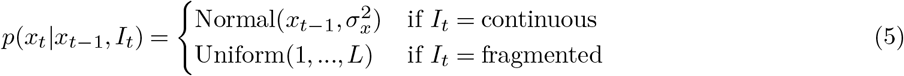

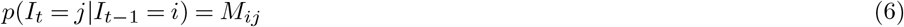

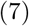

The latent variable is discretized into bins, as is common in the literature (Zhang et al., 1998; Denovellis et al., 2021) and allows for exact posterior computation of the latent. It also gives us the basis function naturally from the eigenvalues *λ*_*b*_ and eigenvectors *v*_*b*_ *∈* ℝ^*L*^ of the GP kernel K evaluated on the bins (gram matrix **K**):

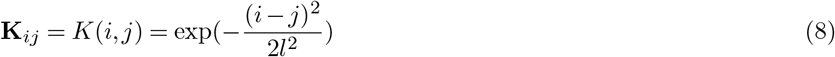

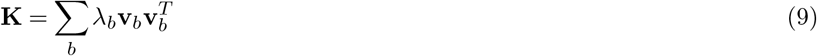

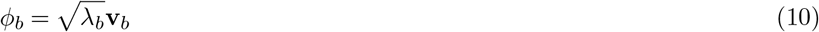

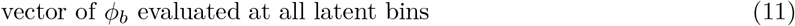

Here we use the radial basis function (RBF) kernel, but our framework is flexible and permits any valid GP kernel and smooth basis functions. We truncate the basis functions where explained variance from the included eigenvalues fall below 99.9%.

### Inference: M-step/Tuning curve estimation

We maximize the log marginal posterior (log likelihood marginalized over latent, conditioned on weights *W* and hyperparameters *θ*, plus log prior):

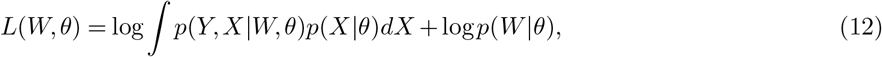

by expectation-maximization (EM) (Dempster et al., 1977), i.e. iterating between 1) maximizing the log complete data posterior (M-step)

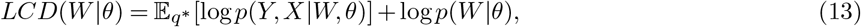

and 2) decoding *q*^*∗*^, i.e. the posterior probability of the latent given the current estimate of *W* (E-step):

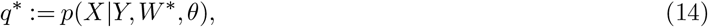

where *W*^*∗*^ is the current estimate of the weights. Crucially, conditioned on *W*, **y**_*t*_ (num. neuron) only depends on *x*_*t*_ and is independent from other times:

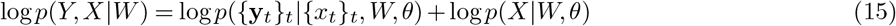

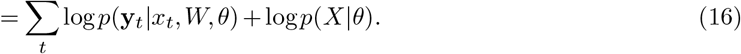

We plug 16 in Eq. 13 and drop the term that does not depend on *W*, we get:

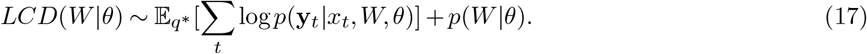

By linearity of expectation and plugging in eq. 14, this equals:

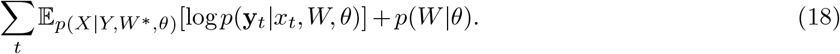

Since *x*_*t*_ only affects log *p*(**y**_*t*_|*x*_*t*_, *W, θ*), taking expectation with respect to all possible latent trajectories (E_*p*(*X*|*Y,W*_ ^*∗*^,*θ*)) for each time bin *t* reduces to taking expectation with respect to possible latent bins at that time 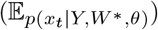. Thus, the objective function reduces to

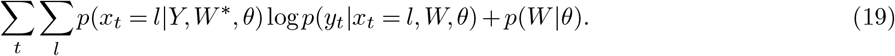

We can then use gradient-based optimizers (or analytic solution if the noise model is Gaussian) to find the optimal *W*^*∗*^ for each M-step iteration. The M-step is very similar to estimating the tuning curves with respect to a given label (e.g. position) using GLM (Hardcastle et al., 2017; Denovellis et al., 2021). The only differences are: 1) *X* here is decoded using the estimate *W*^*∗*^ from the previous iteration and not given. 2) Each time does not correspond to a single latent bin but a distribution, and thus the samples are weighted by the posterior of *X*.

### Trick to speed up M-step

When the number of time bins *T* is large, obtaining gradient from the full dataset, which requires *O*(*T*), can become not only inefficient but even infeasible. Thus, we utilize another trick – rewriting the objective using the summary statistics for each latent bin, another benefit brought by the discretization of the latent. Exchange the order of summation in Eq. 19 (ignoring the prior term for *W* in the manipulation below):

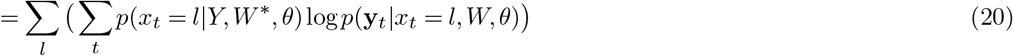

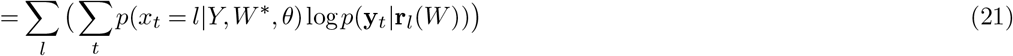

the effect of *W* is through a per latent bin (*l*) rate **r**_*l*_ := *{r*_*ln*_*}*_*n*_, *n* for neuron

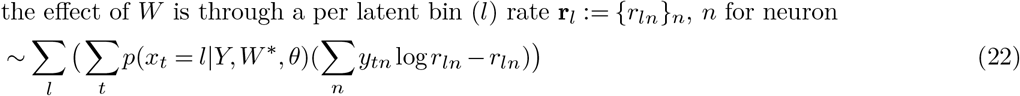

with Poisson observation

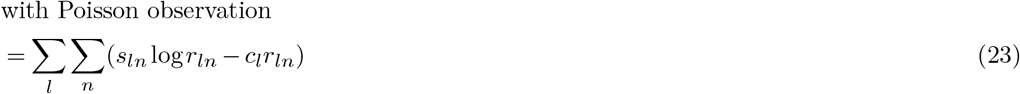

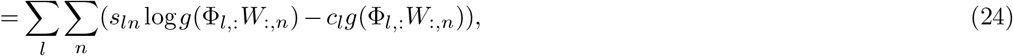

where *c*_*l*_ := ∑_*t*_*p*(*x*_*t*_ = *l* |*Y, W*^*∗*^, *θ*) is the effective occupancy in latent bin *l* and *s*_*ln*_ := ∑_*t*_*p*(*x*_*t*_ = *l* |*Y, W*^*∗*^, *θ*)*y*_*tn*_ is the effective spike count emitted by neuron *n* in latent bin *l*. Φ_*l*,:_ is the *l*-th row of the basis matrix, corresponding to basis functions evaluated at latent bin *l*. Now computing the gradient of *W* is *O*(*L*) and *L << T* . We use ADAM optimizer (Kingma, 2014) with Poisson observation.

### Inference: E-step/State-space decoding

Once we obtain the *W*^*∗*^ from the M-step, we perform state-space decoding using the Bayesian smoother in the same manner as in Denovellis et al. (2021). We first perform Bayesian filtering to recursively get the causal posterior of both the latent and dynamics (omitting the *W* and *θ* in the condition):

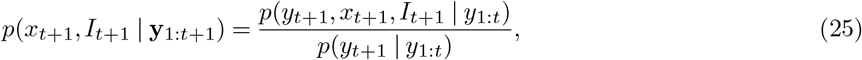

where the numerator is:

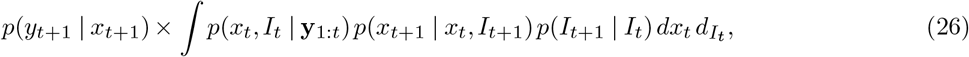

and the denominator is the normalization factor, obtained by marginalizing the numerator over *x*_*t*+1_ and *I*_*t*+1_:

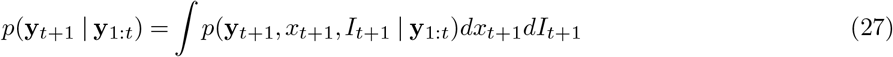

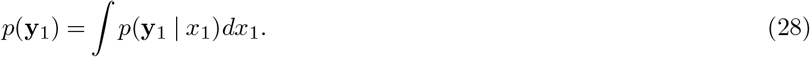

This gives a recursive formula for computing the marginal likelihood. The recursion starts with *p*(*x*_0_, *I*_0_) being a uniform distribution.

Notice *p*(**y**_*t*+1_ | *x*_*t*+1_) is the likelihood of observing the spikes given the latent at time *t*, and thus depends on *W*^*∗*^. (For simplicity of the notation we use integral instead of sum even though *I* is a discrete variable. *x* is a discretized continuous variable).

We then recursively perform Bayesian smoothing for the acausal posterior, starting from the result of the causal posterior *p*(*x*_*T*_, *I*_*T*_ |**y**_1:*T*_) and backward in time:

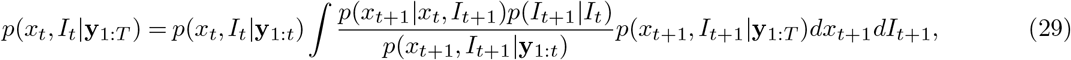

where the causal predictive density is:

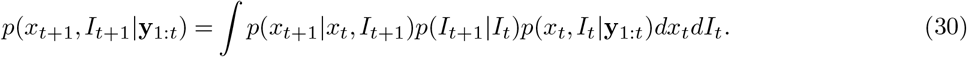

Finally we marginalize over the dynamics to get the posterior estimate of the latent:

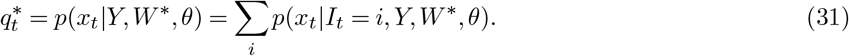

In principle the latent manifold could be any dimension. For computational and analytical reasons we chose to stick to a one-dimensional latent manifold, discretized to 100 bins. While 2-3 dimensions are doable, higher dimensions would lead to an exponential explosion of the number of latent bins. We also found the heatmap of the posterior of latent in time a helpful visualization tool (e.g. Fig. 2), but if the latent dimension is more than one, the visualization would have more complications. This choice does not reflect a belief that the latent intrinsic dimension is necessarily 1D.

Thus the EM steps for the latent variable models correspond to the tuning curve estimation and Bayesian decoding commonly adopted by systems neuroscience. Each step never decreases the log marginal likelihood and is guaranteed to converge to a local minima. The parameters of model is the weights *W*, and the hyperparameters *θ* include: weight prior variance 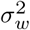, tuning lengthscale *l*, latent transition variance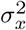, dynamics transition probability matrix *M* . The tuning kernel, transition probability for latent and dynamics can all be changed to custom ones to suit specific needs.

For implementation, all probabilities are represented in log space and marginalization over probabilities are performed using the “log-sum-exp” trick. log *p*(*X, I*| *Y*) is stored as a *T ×*2 *×L* tensor. The filtering and smoothing are performed as matrix summation and logsumexp.

### Initialization

We initialized the posterior *q*^*∗*^ = *p*(*X* | *Y, W, θ*) by independently sampling a positive value for each latent bin and time bin from a uniform distribution. We then re-normalized it to sum to 1 in the latent dimension.

### LVM without Jump

The LVM without jump is the same as the JumpLVM except the model does not contain the dynamics *I*_*t*_, and the prior on the latent transition is the Gaussian random walk as in the “continuous” dynamics of the JumpLVM.

### Simulation

We simulated 10,000 time bins of latent trajectories and dynamics from the generative model with 100 latent bins. We sampled basis function weights to compute tuning curves for 50 neurons and then sampled spikes from the tuning curves evaluated at the latent state in each time bin. The generative tuning parameters and fitted-model tuning parameters were initialized independently. All models were fit for 20 EM iterations. We used tuning lengthscale 10, movement variance 1, P(Jump->Continuous) = 0.01, p(continuous->jump) = 0.01, and prior variance of the weights 1. To test the effect of temporal smoothness in the LVM without jump, we fit it using movement variances 1 and 30. We also simulated continuous-only dynamics using otherwise identical settings and fit these data with JumpLVM (Supplementary Fig. S1B).

After fitting the JumpLVM and LVM without jump on the simulation data, to aid visualization and confirm that JumpLVM correctly identifies the population co-activity patterns, we permuted the latent bins to match the ground truth latent arrangement. For each fitted model, we computed an overlap matrix

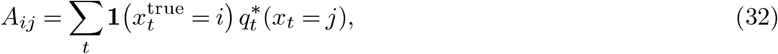

where each entry *A*_*ij*_ measures the total posterior probability assigned to fitted latent bin *j* across all time points at which the true latent was bin *i*. We then found the one-to-one permutation that maximized the summed posterior overlap across all latent bins:

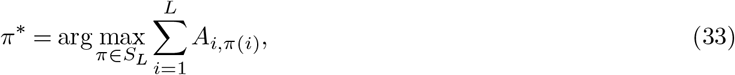

where *S*_*L*_ is the set of all permutations of the *L* latent bins. We applied the same permutation to the latent posterior and tuning curves. This was implemented using `scipy.optimize.linear_sum_assignment.` Because permutation disrupted tuning-curve smoothness, we performed one additional fixed-posterior M-step after permutation for visualization; no additional E-step was performed, and the inferred dynamics were unchanged. This procedure was only applied to the simulated data and not real data.

### Jump clean-up

There are two cases when spurious jumps can happen: 1) when the model does not recover the global latent arrangement (see Supplementary Fig. S1A, rows 1–2 and B, row 2); 2) when the topology of the neural data mismatches that of the latent variable (1D line), because the model has no way of forcing all the similar neural patterns on adjacent latent bins (Supplementary Fig. S2A). Instead of relying on the prior transition kernels, we replaced the prior continuous transition kernel with a tuning-similarity-based kernel and re-decoded to obtain a cleaned jump probability (Supplementary Fig. S2). To minimize the influence from the smoothness in the original learned tuning curves, we first recomputed the empirical tuning curves (no smoothing): for each latent bin, the mean firing rate per neuron under the posterior over the latent from the original decode (posterior-weighted average of spike counts per latent bin). We then defined a Gibbs transition matrix on latent bins: from bin *i* to *j*, the transition weight *G*_*β*_(*i, j*) was given by:

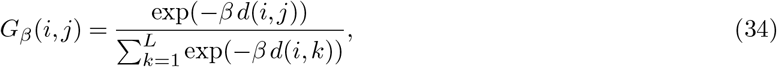

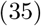

with *d*(*i, j*) the pairwise cosine distance between the tuning vectors of states *i* and *j*, and *β* an inverse-temperature parameter. Rows were normalized to obtain a valid transition matrix. Thus, transitions between similar tuning states are favored and between dissimilar states disfavored. We set *β* by matching this Gibbs transition to a target transition. The target was the latent transition distribution under continuous dynamics only, i.e., the posterior transition from the full model restricted to the continuous regime and renormalized, which reflects how the latent moves when no jump is involved. Only continuous dynamics was used as the target because then the new kernel would be on a similar scale of values as the empirical transition, but we wouldn’t overfit to the actual jumps from the fragmented dynamics. We selected *β* by minimizing, over a discrete grid of *β*s, the KL divergence between the target and the Gibbs transition (row-wise):

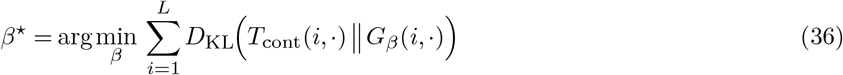

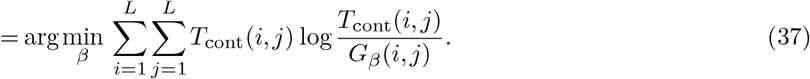

The target *T*_cont_ is the posterior transition kernel under the continuous regime, obtained by aggregating the smoothed two-slice marginal and renormalizing each row:

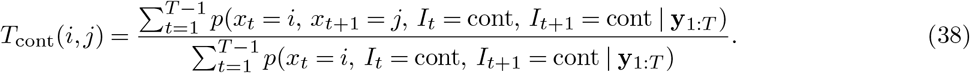

The smoothed two-slice marginal can be obtained from the filtering (Eq.) and smoothing posteriors Eq. 29 (Rabiner, 2002):

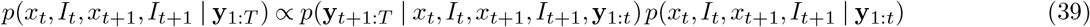

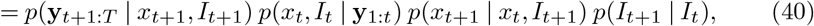

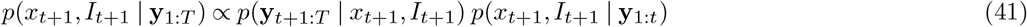

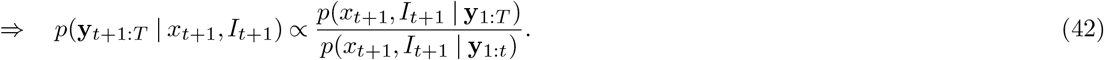

Substituting Eq. 42 into Eq. 40 yields

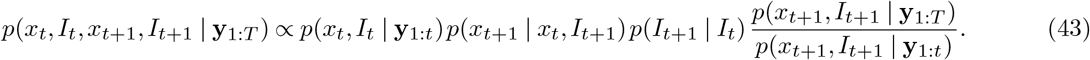

Decoding was then repeated with this fixed “clean” transition kernel to obtain the cleaned posterior over the continuous vs. jump dynamics.

### Jump consistency

One additional safeguard against spurious jumps is to focus only on the ones that occur consistently in different fitted models from different random initializations. We measured the fraction of consistent jumps as a function of: 1) the threshold for binarizing *P* (*Jump*) into jump events vs continuous dynamics; 2) the window to consider two jumps as consistent. Among the five random initializations, we chose the random initialization with the highest model marginal likelihood (marginalizing over latent, conditioned on parameters and hyperparameters), binarized its dynamics posterior *P* (*Jump*), and for each jump event, checked whether 4 out of 5 chains have a jump within the window. We swept the threshold and window size in Fig. 3C, and use threshold= 0.6 and window= 0.5*s* by default.

#### A. Spiking data preprocessing overview

To fit JumpLVM to data, we binned the spike times into 100ms bins (same for the other datasets). The only difference is that the JumpLVM was fitted to the whole session including sleep for the ACh dataset, whereas for T-maze CA1 and the Linear track MEC datasets, we fitted to the Task epoch.

### ACh dataset

Data were collected from adult ChAT-Cre × Ai32 mice (both sexes, 3–6 months), approved by the NYU Langone IACUC and housed on a reverse 12 h light–dark cycle with ad libitum food/water. Mice underwent standard stereotaxic surgeries to deliver opsins/sensors to septal–hippocampal circuits (e.g., cholinergic tools in medial/lateral septum and ACh sensors such as rACh1.4/ACh3.0 in dorsal hippocampus), with optical fibers implanted for medial septum stimulation and hippocampal fiber photometry. Extracellular activity was recorded from CA1/CA3 using multishank silicon probes (64–128 channels), sampled at 20 kHz; target layers were verified online by spiking profiles, ripple expression, and sharp-wave polarity. Behavior comprised home-cage sleep/quiet wakefulness. Fiber-photometry ACh signals were excited (*∼* 561 nm), sampled at 100 Hz, and preprocessed with median filtering, low-/band-pass filtering, double-exponential de-bleaching, and per-session z-scoring. Optogenetic manipulation of medial septum used blue-light activation (473 nm, 20 Hz, 30% duty) and yellow-light inhibition (*∼* 589 nm, constant 5 s), with power at the brain fiber tip calibrated to *∼* 5–10 mW. The session ID of the data we used in the paper are: CIRP005_027_240817_160055 (without stimulation) and CIRP004_029_240827_055134 (with stimulation). Sleep state scoring was performed similarly as for the T-maze dataset.

### T-maze CA1 dataset

For details on animal surgery, training, recording, data preprocessing, and spike sorting / cell body segmentation, we refer to Huszár et al. (2022). In brief, we used the chronic silicon probe recordings from hippocampal CA1 region. The animals were trained on a spatial alternation task on a figure-eight maze. Animals were water restricted before the start of experiments and familiarized to a customized 79×79cm^2^ figure-eight maze raised 61cm above the ground. Over several days after the start of water deprivation, animals were shaped to visit alternate arms between trials to receive a water reward. A 5-s delay in the start area (delay area) was introduced between trials. The position of head-mounted red LEDs (light-emitting diodes) was tracked with an overhead camera at a frame rate of 30Hz. Animals were required to run at least ten trials along each arm (at least twenty trials total) within each session. In all sessions that included maze behavior, animals spent *∼* 120min in the homecage before running on the maze and another *∼* 120min in the homecage afterward for sleep recordings. All behavioral sessions were performed in the mornings (start of the dark cycle). The session id is `e13_26m1_210913.` Sharp wave ripple (SPW-R) and sleep state scoring are provided in the dataset (Huszár et al., 2022). For SPW-R detection (Tingley and Buzsáki, 2020), channels located in the center of the CA1 pyramidal layer were manually selected (https://github.com/buzsakilab/buzcode/blob/master/detectors/detectEvents/bz_FindRipples.m). Broadband LFP was bandpass filtered between 130 and 200Hz using a fourth-order Chebyshev filter, and the normalized squared signal was calculated. SPW-R maxima were detected by thresholding the normalized squared signal at 5s.d. above the mean, and the surrounding SPW-R start–stop times were identified as crossings of 2s.d. around this peak. SPW-R duration limits were set to be between 20 and 200ms. An exclusion criterion was provided by designating a ‘noise’ channel (no detectable SPW-Rs in the LFP) and events detected on this channel were interpreted as false positives (for example, EMG artifacts).

Sleep state scoring was performed as described previously (https://github.com/buzsakilab/buzcode/blob/dev/detectors/detectStates/SleepScoreMaster/SleepScoreMaster.m; see also Watson et al. 2016; Levenstein et al. 2019). First, the LFP was extracted from wideband data by low-pass filtering (sinc filter with a 450-Hz cut-off band) and downsampling to 1,250Hz. Three signals were used for state scoring: broad-band LFP, narrowband theta frequency LFP and electromyogram (EMG). Spectrograms were computed from broadband LFP with fast-Fourier transform in 10-s sliding windows (at 1s) and principal component analysis (PCA) was computed after a z transform. The first PC reflected power in the low (<20-Hz) frequency range, with oppositely weighted power at higher (>32-Hz) frequencies. Theta dominance was quantified as the ratio of powers in the 5-to 10-Hz and 2-to 16-Hz frequency bands. EMG was estimated as the zero-lag correlation between 300- and 600-Hz-filtered signals across recording sites. Soft sticky thresholds on these metrics were used to identify states. Briefly, high LFP PC1 and low EMG were taken to be NREM, high theta and low EMG were considered to be REM and the remaining data were taken to reflect the waking state. All assignments were inspected visually and manually curated wherever appropriate (https://github.com/buzsakilab/buzcode/blob/dev/GUITools/TheStateEditor/TheStateEditor.m). From the x,y trajectories from the head camera tracking, we computed velocity: *v*_*x*_ = *x*^*′*^, *v*_*y*_ = *y*^*′*^ (both smoothed with a gaussian kernel with 0.3s std), speed 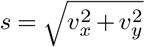 and acceleration: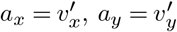.

### Supervised spatial tuning and decoding for T-maze CA1 dataset

For Fig.3B, the baseline is a generalized linear model (Pillow et al., 2008; Hardcastle et al., 2017) using 2D position as the regressor. The GLM uses 2D B-spline basis functions to transform the positions into a design matrix, and uses a soft-plus nonlinearity and Poisson noise model:

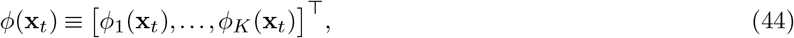

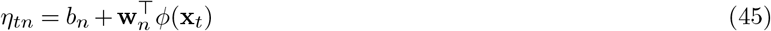

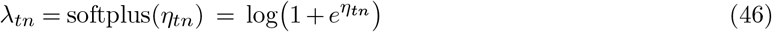

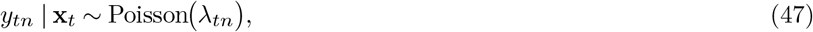

where **x**_*t*_ *∈* ℝ^2^ is the 2D position at time *t* with components 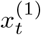 and 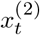 ; *y*_*tn*_ *∈* N_0_ is the spike count of neuron *n* in time bin *t* (for *t* = 1:*T, n* = 1:*N*); *ϕ*(**x**_*t*_) is the output of the *K* basis function; *b*_*n*_*∈* ℝ and **w**_*n*_*∈* ℝ^*K*^ are the intercept and weights for neuron *n*; *λ*_*tn*_ and *y*_*tn*_ are the rate and spike counts for neuron *n* at time *t*. The fitting is via L-BFGS. The ridge regularization strength on **w**_*n*_ and the number of basis functions on is selected via cross validation. The same train-test split is performed on the data for the GLM and the JumpLVM. We simply use the first 80% of the task epoch as the training set (without speed filtering) and the last 20% as the test set. Log_2_-likelihood-per-spike is computed as: log likelihood / (# of spikes) *×* log 2. (Log marginal likelihood was used for the JumpLVM.) Log_2_-likelihood-per-spike gains are obtained by subtracting the log_2_-likelihood-per-spike of an inhomogenous Poisson model, assuming a constant firing rate within the selected epoch. The relative log_2_-likelihood-per-spike gain is the log_2_-likelihood-per-spike gain of the JumpLVM (or LVM without jump) divided by the log_2_-likelihood-per-spike gain of the GLM. The GLM fitting uses the Nemos package (Balzani et al., 2025). In Fig.3B, immobility was time when speed *≤* 2cm/s, and movement was when speed *≥* 5cm/s, a common threshold used in the literature, slightly different from the more fine-grained behavior classification used in the rest of the paper.

For Fig. 3G, the supervised baselines are based on the encoding and decoding models in Denovellis et al. (2021) and implemented using the “replay_trajectory_classification” library (https://doi.org/10.5281/zenodo.13712700). Tuning to the linearized position (binned at 3cm) was fitted via Kernel density estimators (KDE, with a standard deviation of 1 selected from cross validation) (Denovellis et al., 2021). Tuning was fitted only during movement (speed *≥* 5cm/s, as is commonly done for spatial decoding. The track linearization here is based on the linearization algorithm (Loren Frank Lab, 2024), which uses a track graph and projects different 2D positions to different linearized positions (whereas in the linearization in Fig. 3A, from Huszár et al. (2022), the two turns share the same range of linearized positions). State space decoding is performed similar to the JumpLVM, see eq. 26 and 29 (except the tuning curve is constructed using spatial label here). The movement variance was determined to be 1000 via cross validation and the transition probability of the dynamics types are fixed as the default (*P* (stay) = 0.98,*P* (switch) = 0.02. To compare likelihood of the spikes given the predicted firing rates, we form firing rate prediction for the JumpLVM using the average of the tuning over the posterior of the latents:

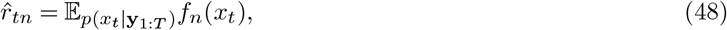

where *f*_*n*_(*x*_*t*_) is the latent tuning curve for neuron *n* evaluated at latent at time *t*. For the supervised model, we form spike prediction from either the spatial tuning curves evaluated at the position labels (“Labeled position LL”), or a posterior average of the tuning curve evaluated at each possible position (“Decoded position LL”, similar to the JumpLVM). The latter measure considers the deviation of the population spatial representation from the position labels, and thus considers correlation in the population to a degree. It serves as a stronger baseline for the JumpLVM to compare against.

For Fig. 4D and Fig. 5B, the supervised encoding and decoding models are the same as in Denovellis et al. (2021), but we used 2-D position instead of 1-D linearized position as the labels. The tuning was also multiplied by the same gain that was selected for JumpLVM. The movement variance was 10 and the standard deviation for the KDE for tuning was 3.

### Behavior type classification

For the T-maze dataset, the latent activation map motivated us to classify the behavior into immobility, off-maze and locomotion.Immobility was defined as epochs with speed *≤* 2.5cm/s. Off-maze was identified by a distance-to-maze criterion and then filtered. Sample positions from the maze was obtained by linearizing 2D position onto T-maze track graph using the track linearization algorithm as before (Loren Frank Lab, 2024). Distance-to-maze was defined as the minimum distance from animal’s position to the maze samples. Candidate off-maze windows were found by finding time bins whose distance-to-maze was *≥* 5cm, then extending them both sides to the first time bins when the distance-to-maze fell below 1cm. It has to further satisfy these criteria: (1) contained at least one time with |angular velocity| *≥* 6 rad/s (angular velocity from 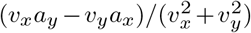, 0.1 s smoothing), and (2) had start-to-end displacement *≤* 15cm (Monaco et al., 2014; Zheng et al., 2024). Locomotion was defined as epochs with speed *≥* 5cm/s.

### Latent classification

For the T-maze dataset, based on the latent activation during different behavior types, we classified latent bins similarly into: locomotion, immobility and off-maze. For a given latent bin, we defined its occurrences as all the time points when it was the maximum a posteriori (MAP) latent. We then checked the number of occurences when the speed of the animal was greater than 5cm/s. The latent was classified as “immobility” latent if that number of time points is below 10 (1s). If it was not an immobility latent, we then checked how often it occurred during off-maze windows. If the off-maze occurrences is greater than 10 bins (1s), then it is classified as an “off-maze” latent. The rest of the latents are “locomotion” latents. This classification makes sure the immobility latent rarely occur during other behavior types, while the off-maze latent could potentially occur during other behavior types.

During hyperparameter sweep (Supplementary Fig. S6), we performed another latent classification to confirm our conclusion did not arise from the particularity of latent classification. we computed the mean posterior mass of latent *k* within each behavior type *b ∈ {*immobility, locomotion, off-maze*}*:

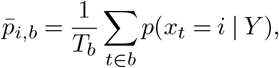

where *T*_*b*_ is the number of time bins assigned to behavior type *b*. We then normalized across behavior types for each latent to obtain a behavior score:

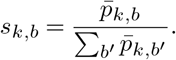

For instance, if a latent occurs in both locomotion and off-maze, but out of a total of 10 time bins from off-maze it occured 9 times, and out of a total of 1000 time bins from locomotion it also occured 9 times, it would have a higher score for off-maze. We then classsified based on a category-specific threshold (e.g. we want immobility latent to occur more strictly during immobility, whereas locomotion could occur during immobility during remote representations): immobility: *≥* 0.8, off-maze and locomotion: *≥* 0.6.

### Population burst event (PBE) definition

We adapt the population burst event (PBE) definition from Yang et al. (2024). First, spikes from the population were binned by 2ms and smoothed with a Gaussian filter (7.5ms standard deviation), and Z-scored. Second, windows where the population spike counts exceed 3 standard deviations were kept and extended in both directions to when the spike counts fall back to the mean. Third, windows below 50ms and above 500ms were discarded. These define the windows of population burst events (PBEs). Only putative pyramidal cells are included in all analyses. When JumpLVM and supervised spatial decoder were applied to the PBEs, spikes within PBEs were binned into 20ms bins.

### Reactivation analysis with JumpLVM

For the T-maze dataset, After JumpLVM was fit to the Task epoch, we performed Naive Bayes decoding on the spike counts during PBEs to determine reactivation (without dynamics):

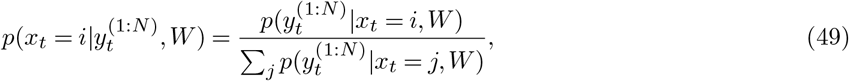

which also gives the marginal likelihood of each event:

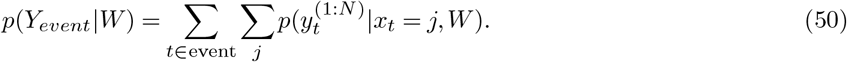

The likelihood 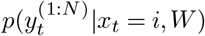 depends on the tuning curves learned from the Task epoch. To account for the potential change in firing statistics during PBEs, we multiplied the tuning curves by a gain factor greater than 1. The gain factor was chosen to maximize the fraction of significant events (same shuffle tests as in sec.) among half of the events (randomly selected).

We used two shuffle controls to build null distributions for the log marginal likelihood. Neuron-ID shuffling permutes the mapping between spike trains and neuron identities while keeping the same set of tuning curves, testing whether high likelihood requires the correct neuron–tuning correspondence. Independent circular shifting rotates each neuron’s spike train by a random offset, preserving per-neuron firing statistics but destroying across-neuron co-activity, testing whether likelihood gains reflect coordinated ensemble structure rather than rate alone. If an event’s marginal likelihood exceeds the 95-percentile in both shuffles, we call it “significant.” All the subsequent analysis of reactivation and replay was restricted to the significant PBEs. This procedure was the same when we used the spatial tuning curves (see) to determine significant reactivation.

### Replay analysis with JumpLVM

To study the temporal dynamics of reactivation, we updated the fitted JumpLVM to incorporate learned latent dynamics within different behavior types, and perform state-space decoding of significant PBEs to decode the new dynamics types. We finally classified each PBE by the dominant dynamics type.

#### Compute behavior-specific transition dynamics

The fitted latent variable model was used to decode the full session to obtain the posterior marginal over latent bins *p*(*x*_*t*_ | *y*_1:*T*_). For each behavior type *b* (e.g., immobility, offmaze, locomotion), we restricted the decoded posterior to the corresponding behavioral intervals. Within each interval we treated time as a single “episode” (no transitions across interval boundaries). We computed expected latent transition counts from the posterior marginals:

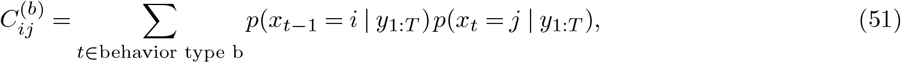

where the sum runs over consecutive time bins within episodes. The behavior-specific transition matrix was then the row-normalized count matrix:

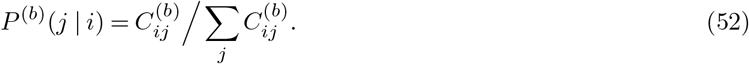

This yielded one *L×L* transition matrix per behavior type (*L* = number of latent states), denoted *{P* ^(*b*)^*}*.

#### Multi-dynamics decoding on PBEs

These transition probabilities replaced the prior continuous transition probability. The final state-space decoder had these three dynamics types plus the fragmented dynamics as described before. The transition probability into a different dynamics was given by (1 *− p*_*stay*_))*/*3, where the stay probability *p*_*stay*_ was fixed as 0.95. We performed the forward-backward smoothing as described before (eq. 29) and obtained the marginal probabilities of the dynamics types at each time. The probabilities were time-averaged within each PBE. The PBE was classified as one behavior type if the corresponding dynamics probability*≥* 0.8. Otherwise it was fragmented or unclassified.

### Calibration analysis for classifying behavior types

In Supplementary Figure. S6, we confirmed whether the latent transitions learned within different behavior types could indeed be used to predict that behavior type on held-out data. If not, then using those dynamics types to classify replay would be meaningless. After fitting the JumpLVM, we splitted the epochs from each behavior type into train and test 10 times. The transition probabilities were fitted only during the train epochs and the posterior for the dynamics types during the test epochs were averaged across time.

### Linear track MEC dataset

The details of the Neuropixel recording of the MEC are given in (Vollan et al., 2025) (https://doi.org/10.25493/R5FR-EDG). Long evans rats were implanted with Neuropixels phase-3A (single-shank) or 2.0 (multi-shank) probes targeting medial entorhinal cortex/parasubiculum and/or hippocampus using standard chronic stereotaxic procedures; implants were secured with dental cement and postoperative analgesia was provided. Neural data were amplified and digitized on-probe and acquired at 30 kHz with SpikeGLX. Head pose was tracked in 3D with an OptiTrack system (retroreflective markers), then projected to the horizontal plane for 2D position and head-direction estimates; data streams (neural, motion capture, video when present) were synchronized via TTL/IR pulse trains. The linear-track task consisted of shuttling on a 200 cm track with liquid rewards at each end; animals were pretrained to perform *∼*40 laps per session, and recording sessions lasted *∼*45–66 min with *∼*18–48 min of running between rewards. The session we used is under the folder: sharing_v4/navigation/lt/26648_1.

## Supplementary Information

**Figure S1.**
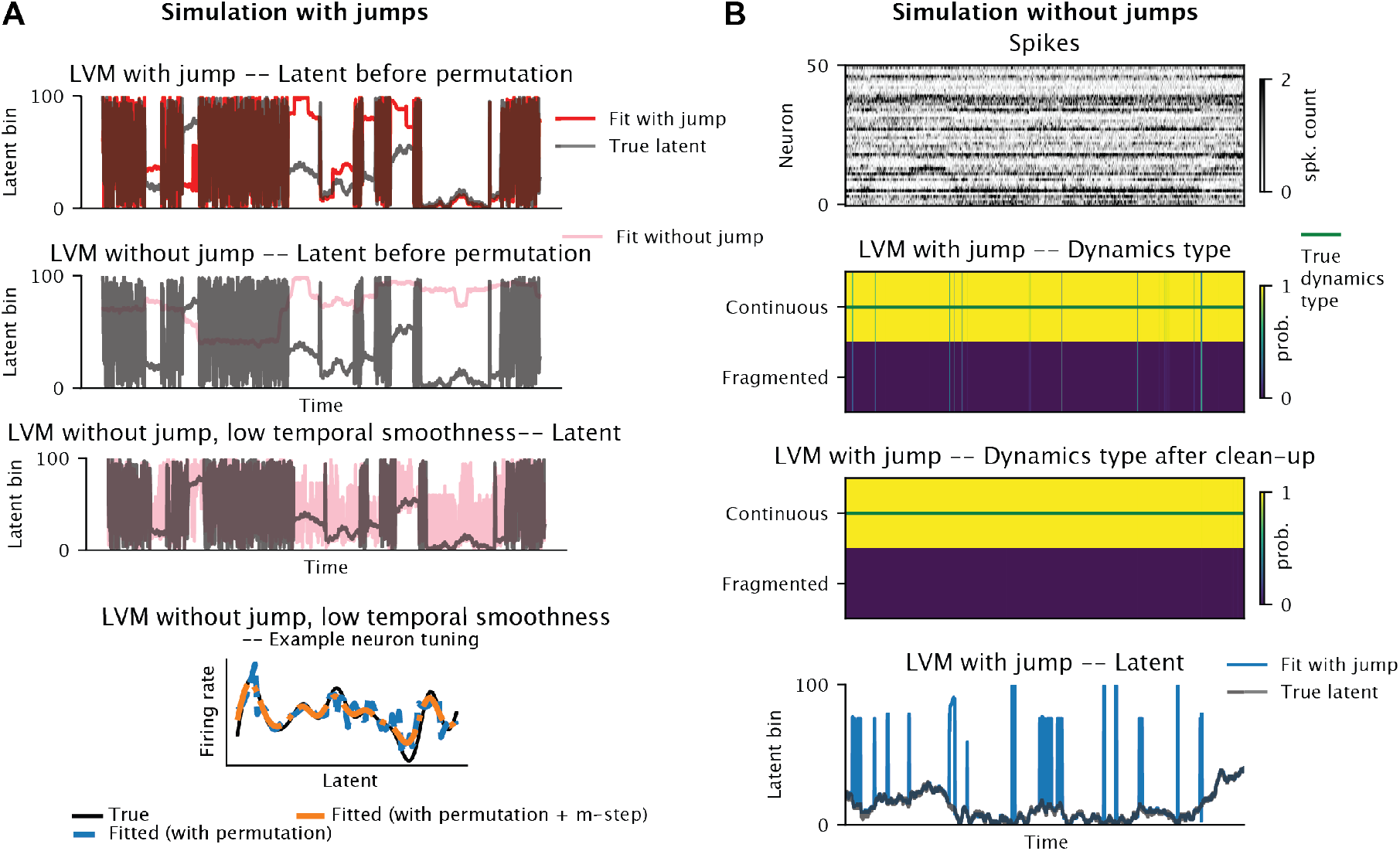
Controls for jump-containing and continuous-only simulations. **(A)** Additional results on the simulation data with jump as in Fig. 1. Row 1: similar to Fig. 1E top, maximum a posteriori (MAP) latent bin in an example time window, before the latent permutation. Row 2: similar to row 1, but the inferred latent is from the LVM without jump as in Fig. 1E bottom. Row 3: similar to row 2, but the inferred latent is from the LVM without jump and lower temporal smoothness (higher movement variance). This panel together with Fig. 1E bottom shows that the LVM without jump struggles when the dynamics types include both continuous and fragmented dynamics, at both low and high movement variances (1 and 30). Row 4: similar to Fig. 1F right, but from the LVM without jump and lower temporal smoothness (higher movement variance). **(B)** Row 1: Raster of spikes generated from a latent variable model without jump. Row 2: Learned JumpLVM mostly decodes continuous dynamics, with occasional false positive jumps. Row 3: After the clean-up step demonstrated in Supplementary Fig. S2, the false positive jumps are removed. Row 4: Learned latent mostly matches the ground truth. Some of the apparent jumps are caused by latent permutation.

**Figure S2.**
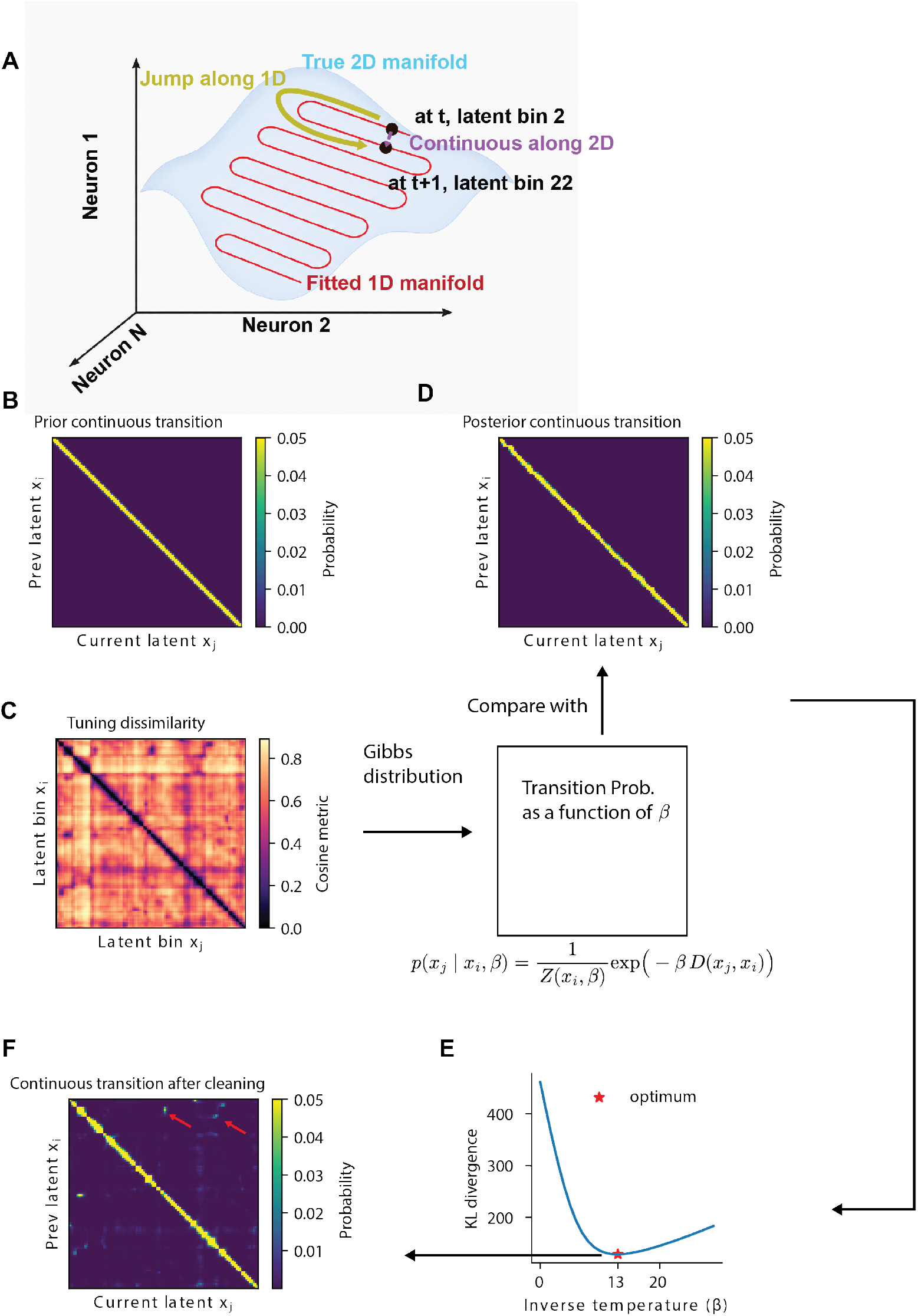
Schematics for cleaning up spurious jumps. **(A)** Schematic for why a clean-up step might be needed, in addition to the failure mode demonstrated in Supplementary Fig. S1B, row 2. If the true manifold (blue) is higher than 1D, then fitting a 1D manifold (red) could allocate points (black dots) that are close-by on the true manifold to distant latent bins, thus mislabeling a small step (purple) as a jump (yellow). **(B)** Prior continuous transition. **(C)** Left: tuning dissimilarity matrix, each entry is the cosine metric between the population tuning corresponding to a pair of latent bins. Right: The tuning dissimilarity matrix is turned into a transition probability using the Gibbs distribution. The matrix is a function of the inverse temperature parameter *β* that needs to be selected. To minimize the potential bias of smoothness on tuning dissimilarity, we recomputed the tuning here using the posterior weighted spike count and occupancy, without smooth basis functions. **(D)** To get a sensible *β*, we compared the transition probability derived from tuning dissimilarity in (C) with the posterior continuous transition probability (the state-space decoder can output this as a byproduct of the forward-backward smoothing, just like in HMM). We only compared with the continuous state because, had we compared with the full posterior transition probability, we would risk including genuine jumps under the “continuous” dynamics. (**E**) The *β* that minimizes the KL divergence (posterior continuous transition vs tuning dissimilarity based) (summed across rows). (**F**) The transition probability that was used for the continuous dynamics after the clean-up, derived from the optimal *β* from (E). Notice the off diagonal transitions indicated by the arrows.

**Figure S3.**
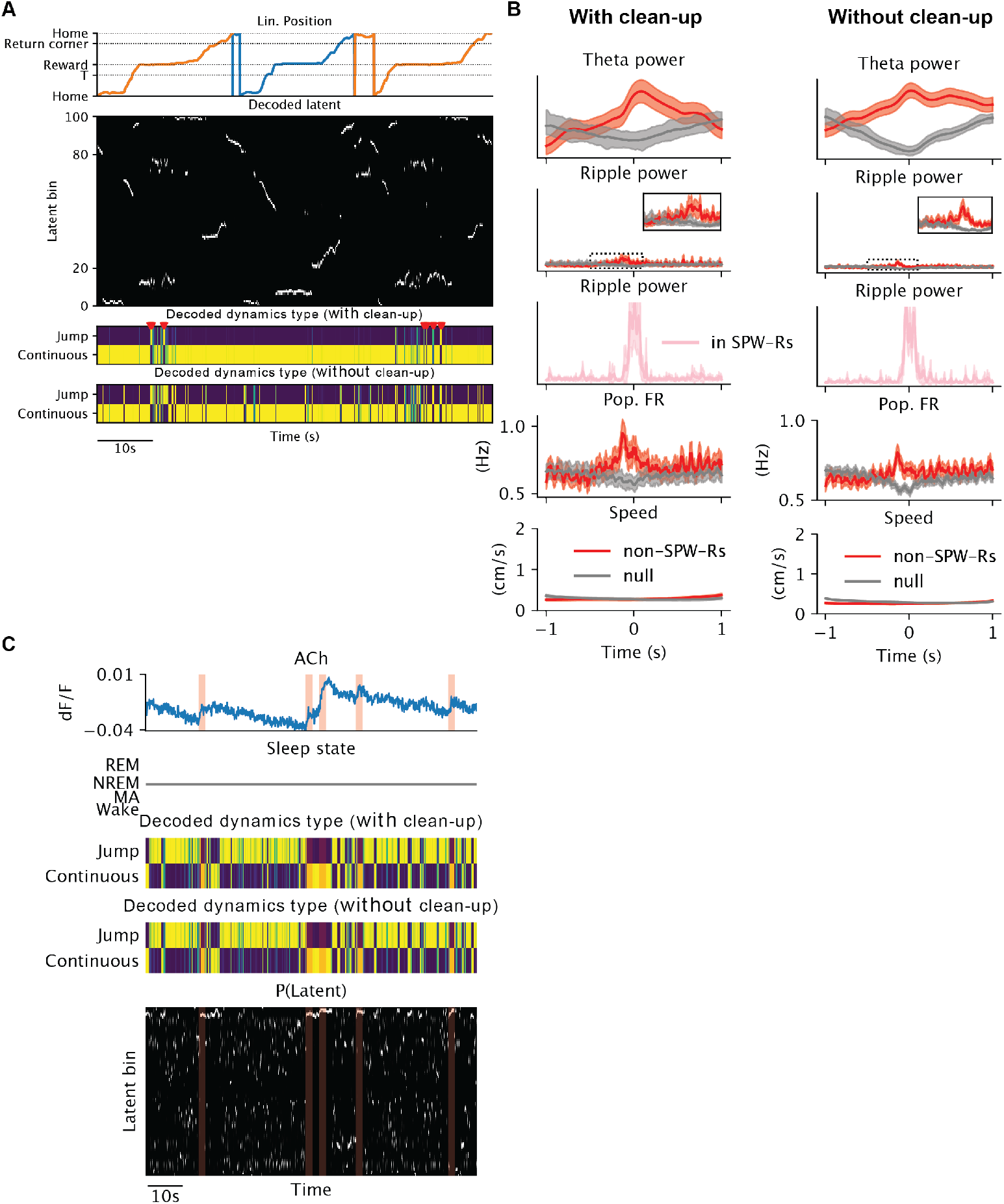
Effects of clean-up. **(A)** Demonstration for how the clean-up step affects the decoded dynamics type for the T-maze data, for some example trials. Similar to Fig. 3A. Row 1: Linearized position in time, colored by turns. Row 2: Posterior probability of the latent. Brighter colors means higher probability. Row 3: Posterior probability of the dynamics type with the clean-up step. Yellow means higher probability. Red markers mark the jump events after the clean-up. Row 4: similar to row 3, but without the clean-up step. Note row 3 has fewer jumps compared to row 4. **(B)** Peri-jump quantities similar to Fig. 3F, using jump events extracted from the decoded dynamics types with (left) vs without (right) the clean-up step. The two columns do not differ in qualitative ways. **(C)** Similar to Fig.2C, with the additional row 3 showing the decoded dynamics type without the clean-up step. Top to bottom: ACh fluoresence, sleep state, posterior probability of the dynamics type after the clean-up step, probability of the dynamics type before the clean-up step, posterior probability of the latent bin. Shaded boxes mark 5s windows after the detected onsets of ACh ramps that occur during NREM.

**Figure S4.**
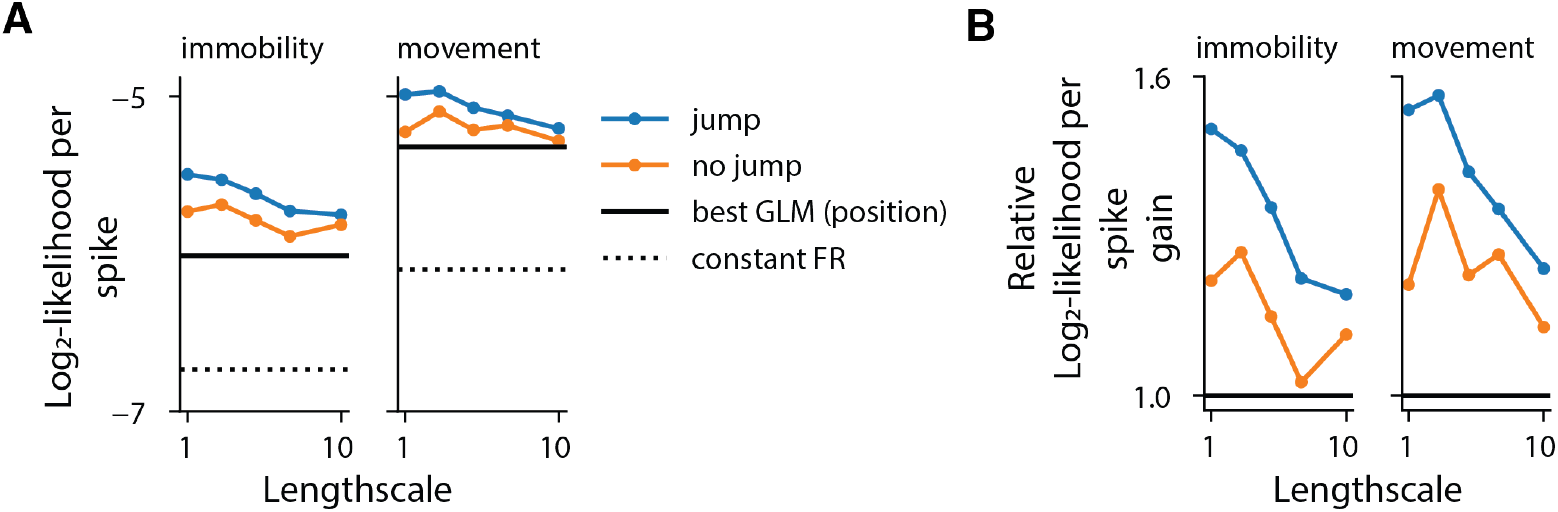
JumpLVM model comparison using a 20ms bin size. **(A)** Log_2_-likelihood-per-spike of different models applied to the T-maze dataset as in Fig.3B, but the spikes are now binned with 20ms bins. **(B)** Same data as in (A), but zoomed in to show the relative log_2_-likelihood-per-spike gain. The gain is with respect to an inhomogeneous Poisson (i.e. constant firing rate) baseline model. The gain is then normalized such that the log_2_-likelihood-per-spike gain of the best GLM with a position regressor is 1.

**Figure S5.**
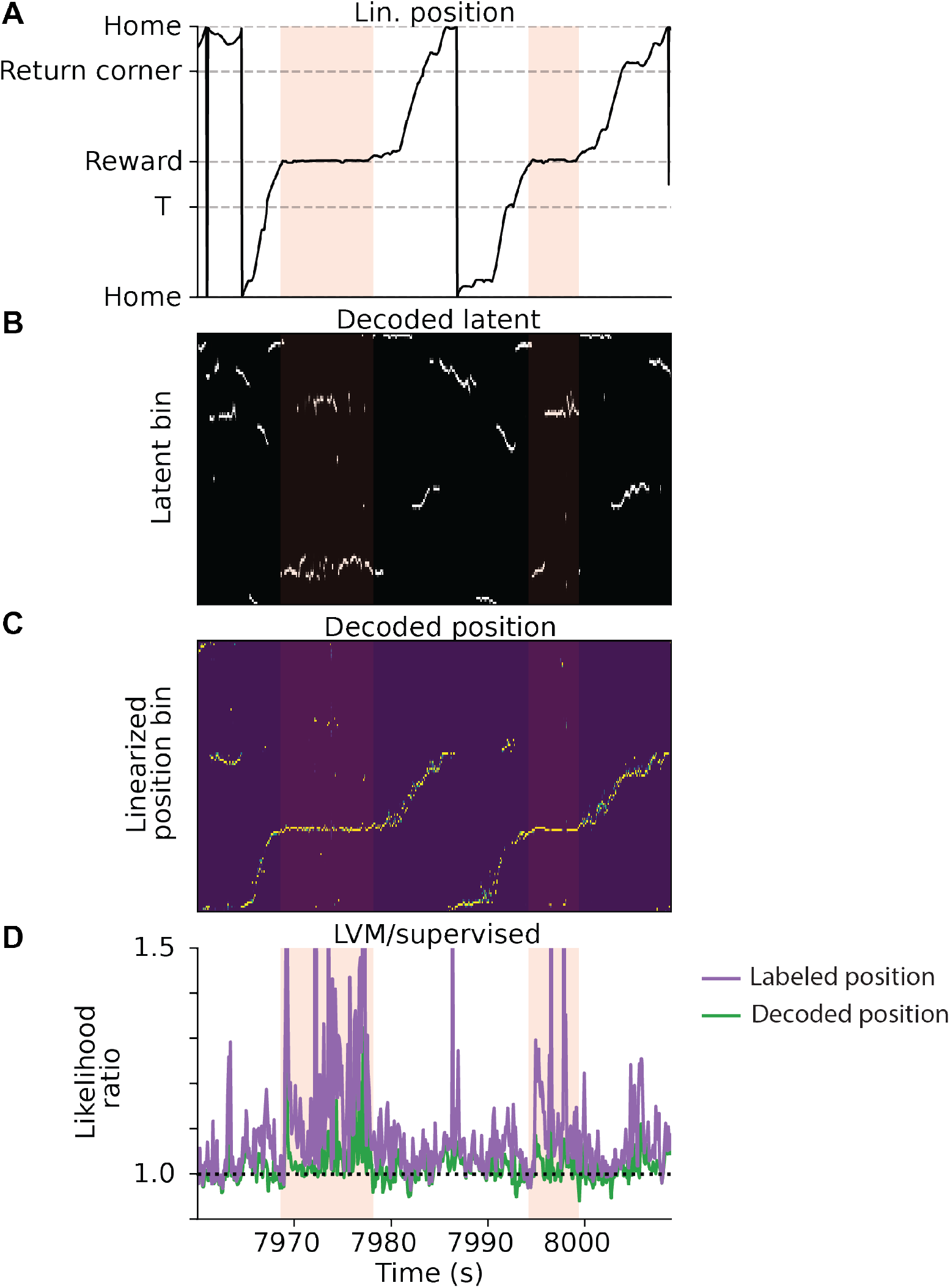
Supervised spatial decoding failed to capture latent difference on an error trial. Plots similar to Fig. 3G. Zoomed in window containing a correct and the subsequent error trial visiting the same arm, as highlighted in Fig. 3A. **(A)** Linearized position. **(B)** Posterior probability of the latent. Jump probability in red line. **(C)** Posterior probability of the decoded position from supervised spatial decoding. **(D)** Likelihood ratio between the JumpLVM and supervised models. The likelihood for each model is computed from the expected firing rates and the Poisson observation. For the labeled position (purple), expected firing rates are computed by evaluating the tuning curves at the given position label. For the decoded position (green) and the JumpLVM, expected firing is the posterior weighted average of the tuning curves evaluated at each position / latent bin.

**Figure S6.**
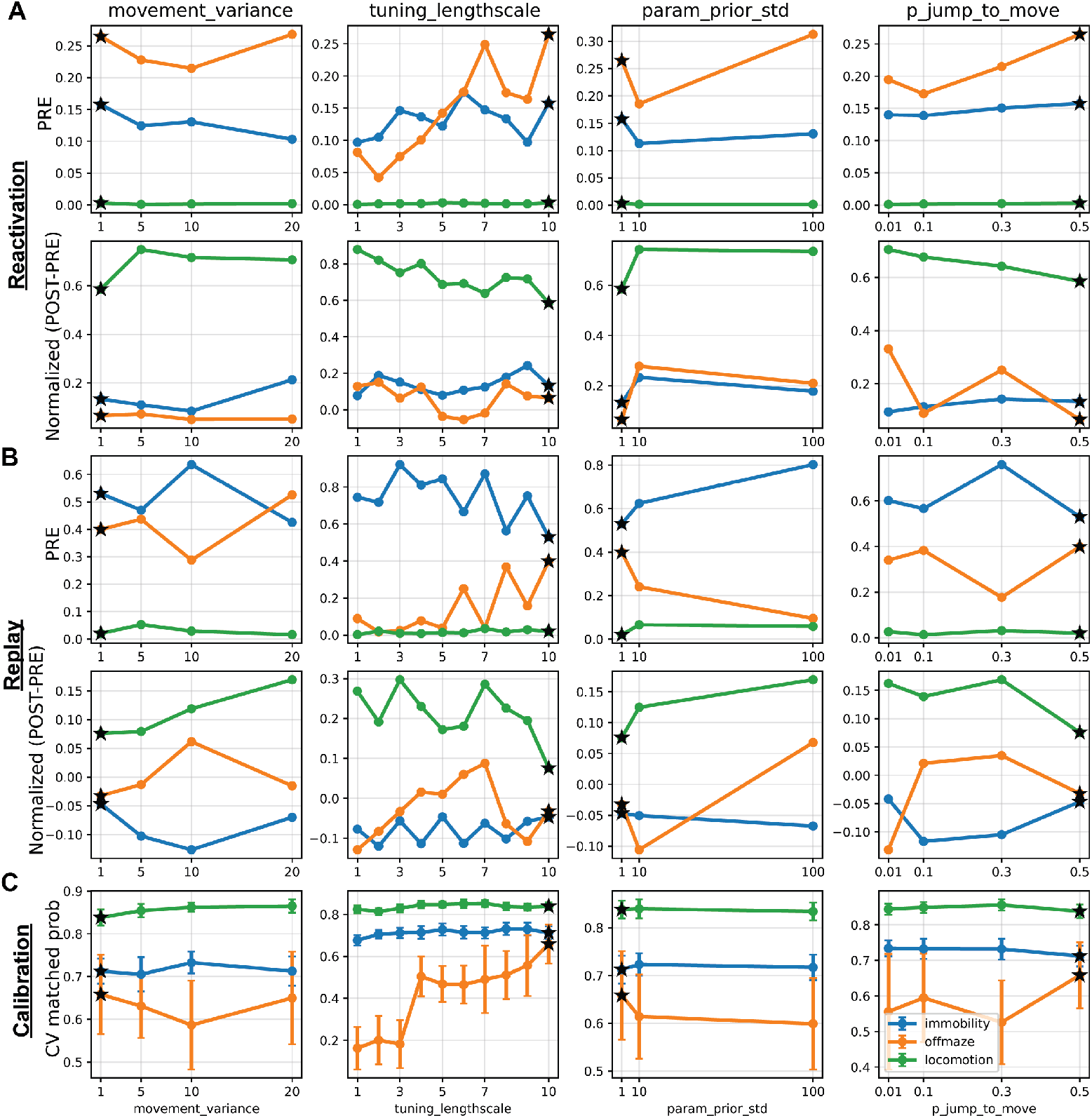
Reactivation and replay analyses are insensitive to hyperparameters. Starting from the default set of hyperparameters used in the main text (marked by black stars), we swept each hyperparameter to observe its effect on the reactivation and replay analysis. Each column corresponds to a hyperparameter being swept, and X axes are the values of the hyperparameter being swept. (**A**) Reactivation analysis similar to Fig. 4E. Each colored line is one category of latent bins. Top row: posterior probability of the latent from one behavior type (max within significant NREM PBE, mean across PBEs, mean across latent bins; for behavior type classification, see *Methods*). Bottom row: measure of reactivation: (POST-PRE)/(POST+PRE), averaged over latent bins. The result is not sensitive to the hyperparameters in this range. **(B)** Replay analysis similar to Fig. 5D. Each line is one dynamics type. Top row: probability of each dynamics type averaged within PBE, median across significant PBEs during the PRE-task NREM sleep. Bottom row: the same aggregation process as the top row to get POST and PRE, but plotting the measure of replay: (POST-PRE)/(POST+PRE). While the result is largely similar, the PRE probability for off-maze is low when the tuning lengthscale is low. **(C)** Calibration analysis of how well the learned latent transition can predict the behavior type on held out data during the task epoch. Each line is a behavior type. Y axis is: for time bins classified as one behavior type, the average probability of the corresponding dynamics type. The error bars mark the 1 standard deviation among 10 cross validation (CV) splits (50-50 train-test). While most hyperparmeters don’t affect this, low tuning lengthscale has poor calibration for the off-maze behavior, justifying the choice of a higher lengthscale.

**Figure S7.**
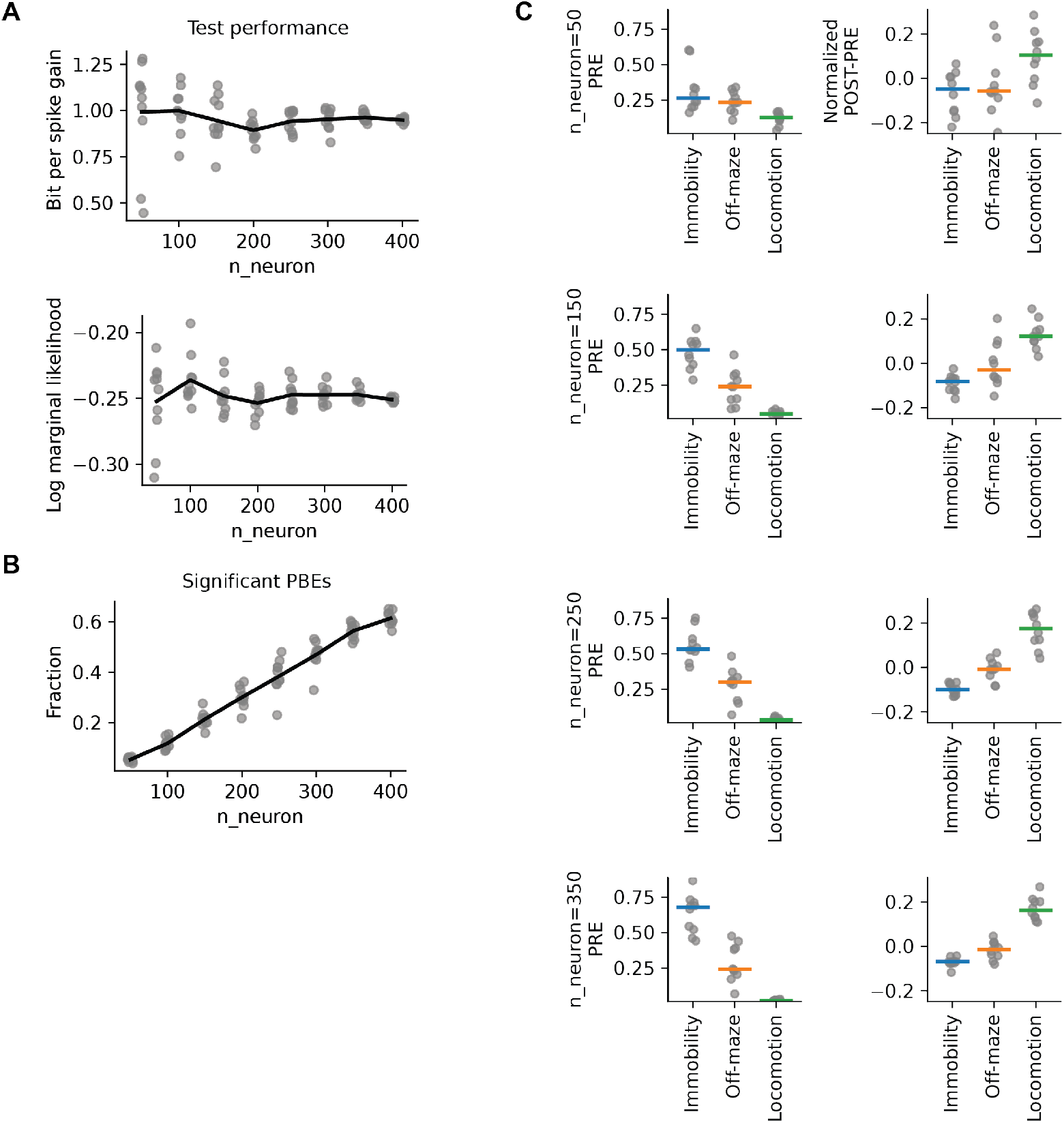
Downsampling of neurons on the T-maze dataset. Each grey dot is from the JumpLVM fitted with one random subset of neurons. At each n_neuron we sampled ten times. **(A)** Held-out data performance as a function of the number of neurons. Performance was measured by bit-per-spike relative to the baseline of inhomogeneous Poisson model (top) or log marginal likelihood per neuron per time bin (bottom). While the performance is similar overall, the variance reduces as more neurons are included. **(B)** The fraction of significant PBEs as a function of the number of neurons. Fewer PBEs are identified as significant when fewer neurons are included. **(C)** Replication of the replay result in Fig. 5D across different numbers of included neurons. Left: each dot is the probability of a given dynamics type averaged within PBE, median across significant PBEs during the PRE-task NREM sleep. Right: similar aggregation process as the left to get POST and PRE, but plotting the measure of replay: (*POST − PRE*)*/*(*POST* + *PRE*).

**Figure S8.**
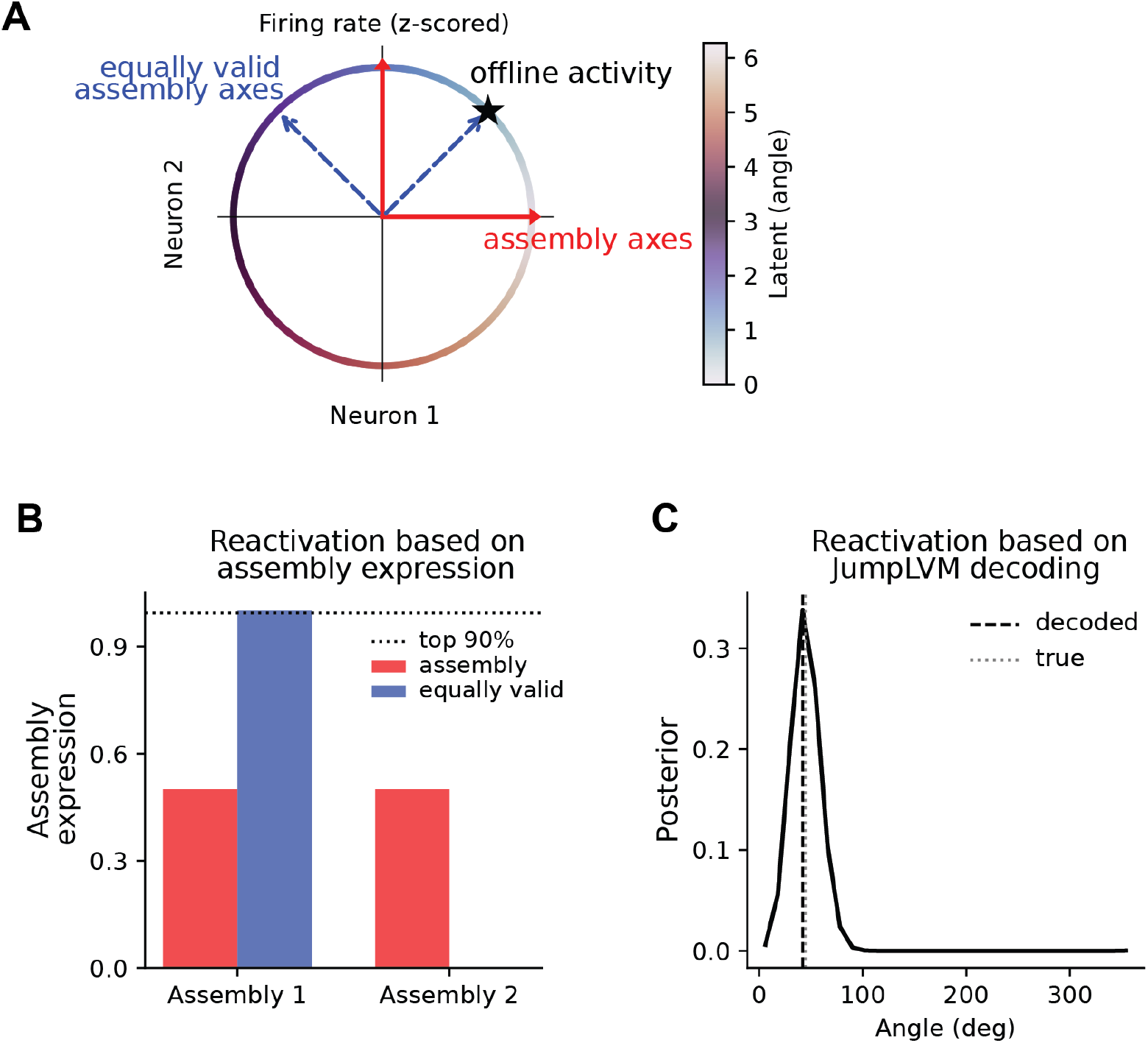
Assembly detection methods fail on non-linear neural manifolds. **(A)** In this toy model, the firing rate vectors of two neurons lie on a circle, colored by the one-dimensional latent variable, for instance the head direction. The red arrows and blue dotted arrows are two sets of equally valid principal component / assembly axes. The black star marks an example firing rate vector that could occur during offline activites. **(B)** The assembly expression strength of the black star firing vector in (A), according to different choices of the assembly axes. The strength is measured as the square of the dot product between the firing vector and the assembly axes. The black dotted line marks the threshold for “reactivation events.” (Assuming the candidate firing vectors during offline is the same circle, we computed the expression strengths and used the 90-percentile as the threshold.) Based on one choice (red) this vector does not count as a reactivation event, while on another choice (blue) it passes the threshold. Since this firing vector is selected from the manifold, it should clearly be considered a reactivation event, thereby showing the arbitrariness of assembly methods when applied to data on a manifold that is nonlinearly embedded. **(C)** JumpLVM correctly decodes the offline firing vector (after aligning the latent space with the true latent).

## Bibliography

Angelaki, D., Benson, B., Benson, J., Birman, D., Bonacchi, N., Bougrova, K., Bruijns, S. A., Carandini, M., Catarino, J. A., et al. (2025). A brain-wide map of neural activity during complex behaviour. Nature, 645(8079):177–191.

Aronov, D., Nevers, R., and Tank, D. W. (2017). Mapping of a non-spatial dimension by the hippocampal–entorhinal circuit. Nature, 543 (7647):719–722.

Averbeck, B. B., Latham, P. E., and Pouget, A. (2006). Neural correlations, population coding and computation. Nature reviews neuroscience, 7(5):358–366.

Balzani, E., Lakshminarasimhan, K., Angelaki, D., and Savin, C. (2020). Efficient estimation of neural tuning during naturalistic behavior. Advances in Neural Information Processing Systems, 33:12604–12614.

Balzani, E., Bagi, B., Biswas, A., Broderick, W. F., Venditto, S. J., Lewis, C., Maura, C., Crespo, P., Viejo, G., Modumudi, P., Thomson, E., and Williams, A. Nemos: Neural models, (2025). URL 10.5281/zenodo.17553287.

Berners-Lee, A., Feng, T., Silva, D., Wu, X., Ambrose, E. R., Pfeiffer, B. E., and Foster, D. J. (2022). Hippocampal replays appear after a single experience and incorporate greater detail with more experience. Neuron, 110(11):1829–1842.

Brown, E. N., Frank, L. M., Tang, D., Quirk, M. C., and Wilson, M. A. (1998). A statistical paradigm for neural spike train decoding applied to position prediction from ensemble firing patterns of rat hippocampal place cells. Journal of Neuroscience, 18(18):7411–7425.

Burke, S. N., Maurer, A. P., Nematollahi, S., Uprety, A. R., Wallace, J. L., and Barnes, C. A. (2011). The influence of objects on place field expression and size in distal hippocampal ca1. Hippocampus, 21 (7):783–801.

Churchland, M. M., Cunningham, J. P., Kaufman, M. T., Foster, J. D., Nuyujukian, P., Ryu, S. I., and Shenoy, K. V. (2012). Neural population dynamics during reaching. Nature, 487(7405):51–56.

Cunningham, J. P. and Yu, B. M. (2014). Dimensionality reduction for large-scale neural recordings. Nature neuroscience, 17(11):1500–1509.

Davidson, T. J., Kloosterman, F., and Wilson, M. A. (2009). Hippocampal replay of extended experience. Neuron, 63(4):497–507.

Dayan, P. and Abbott, L. F. Theoretical neuroscience: computational and mathematical modeling of neural systems. MIT press, (2005).

De, A. and Chaudhuri, R. (2023). Common population codes produce extremely nonlinear neural manifolds. Proceedings of the National Academy of Sciences, 120(39):e2305853120.

Dempster, A. P., Laird, N. M., and Rubin, D. B. (1977). Maximum likelihood from incomplete data via the em algorithm. Journal of the royal statistical society: series B (methodological), 39(1):1–22.

Denovellis, E. L., Gillespie, A. K., Coulter, M. E., Sosa, M., Chung, J. E., Eden, U. T., and Frank, L. M. (2021). Hippocampal replay of experience at real-world speeds. Elife, 10:e64505.

Diba, K. and Buzsáki, G. (2007). Forward and reverse hippocampal place-cell sequences during ripples. Nature neuroscience, 10(10):1241–1242.

Dragoi, G. and Tonegawa, S. (2011). Preplay of future place cell sequences by hippocampal cellular assemblies. Nature, 469(7330):397–401.

Durstewitz, D., Vittoz, N. M., Floresco, S. B., and Seamans, J. K. (2010). Abrupt transitions between prefrontal neural ensemble states accompany behavioral transitions during rule learning. Neuron, 66(3):438–448.

Farooq, U. and Dragoi, G. (2019). Emergence of preconfigured and plastic time-compressed sequences in early postnatal development. Science, 363(6423):168–173.

Foster, D. J. and Wilson, M. A. (2006). Reverse replay of behavioural sequences in hippocampal place cells during the awake state. Nature, 440(7084):680–683.

Gauthier, J. L. and Tank, D. W. (2018). A dedicated population for reward coding in the hippocampus. Neuron, 99(1):179–193.

George, D., Rikhye, R. V., Gothoskar, N., Guntupalli, J. S., Dedieu, A., and Lázaro-Gredilla, M. (2021). Clone-structured graph representations enable flexible learning and vicarious evaluation of cognitive maps. Nature communications, 12(1):2392.

George, T. M., Glaser, P., Stachenfeld, K., Barry, C., and Clopath, C. (2024). Simpl: Scalable and hassle-free optimization of neural representations from behaviour. bioRxiv, pages 2024–11.

Georgopoulos, A. P., Schwartz, A. B., and Kettner, R. E. (1986). Neuronal population coding of movement direction. Science, 233(4771): 1416–1419.

Gillespie, A. K., Maya, D. A. A., Denovellis, E. L., Liu, D. F., Kastner, D. B., Coulter, M. E., Roumis, D. K., Eden, U. T., and Frank, L. M. (2021). Hippocampal replay reflects specific past experiences rather than a plan for subsequent choice. Neuron, 109(19):3149–3163.

Grosmark, A. D. and Buzsáki, G. (2016). Diversity in neural firing dynamics supports both rigid and learned hippocampal sequences. Science, 351(6280):1440–1443.

Gulati, T., Ramanathan, D. S., Wong, C. C., and Ganguly, K. (2014). Reactivation of emergent task-related ensembles during slow-wave sleep after neuroprosthetic learning. Nature neuroscience, 17(8):1107–1113.

Gulati, T., Guo, L., Ramanathan, D. S., Bodepudi, A., and Ganguly, K. (2017). Neural reactivations during sleep determine network credit assignment. Nature neuroscience, 20(9):1277–1284.

Gupta, A. S., Van Der Meer, M. A., Touretzky, D. S., and Redish, A. D. (2010). Hippocampal replay is not a simple function of experience. Neuron, 65(5):695–705.

Gupta, A. S., Van Der Meer, M. A., Touretzky, D. S., and Redish, A. D. (2012). Segmentation of spatial experience by hippocampal theta sequences. Nature neuroscience, 15(7):1032–1039.

Hanks, T. D., Kopec, C. D., Brunton, B. W., Duan, C. A., Erlich, J. C., and Brody, C. D. (2015). Distinct relationships of parietal and pre-frontal cortices to evidence accumulation. Nature, 520(7546):220–223.

Hardcastle, K., Maheswaranathan, N., Ganguli, S., and Giocomo, L. M. (2017). A multiplexed, heterogeneous, and adaptive code for navigation in medial entorhinal cortex. Neuron, 94(2):375–387.

Hollup, S. A., Molden, S., Donnett, J. G., Moser, M.-B., and Moser, E. I. (2001). Accumulation of hippocampal place fields at the goal location in an annular watermaze task. Journal of Neuroscience, 21 (5):1635–1644.

Huszár, R., Zhang, Y., Blockus, H., and Buzsáki, G. (2022). Preconfigured dynamics in the hippocampus are guided by embryonic birthdate and rate of neurogenesis. Nature Neuroscience, 25(9):1201–1212.

Jensen, K., Kao, T.-C., Tripodi, M., and Hennequin, G. (2020). Manifold gplvms for discovering non-euclidean latent structure in neural data. Advances in Neural Information Processing Systems, 33:22580–22592.

Johnson, A. and Redish, A. D. (2007). Neural ensembles in ca3 transiently encode paths forward of the animal at a decision point. Journal of Neuroscience, 27(45):12176–12189.

Johnson, A., Seeland, K., and Redish, A. D. (2005). Reconstruction of the postsubiculum head direction signal from neural ensembles. Hippocampus, 15(1):86–96.

Josselyn, S. A. and Tonegawa, S. (2020). Memory engrams: Recalling the past and imagining the future. Science, 367(6473):eaaw4325.

Karlsson, M. P. and Frank, L. M. (2009). Awake replay of remote experiences in the hippocampus. Nature neuroscience, 12(7):913–918.

Kay, K., Chung, J. E., Sosa, M., Schor, J. S., Karlsson, M. P., Larkin, M. C., Liu, D. F., and Frank, L. M. (2020). Constant sub-second cy-cling between representations of possible futures in the hippocampus. Cell, 180(3):552–567.

Khemakhem, I., Kingma, D., Monti, R., and Hyvarinen, A. Variational autoencoders and nonlinear ica: A unifying framework. In International conference on artificial intelligence and statistics, pages 2207–2217. PMLR, (2020).

Kim, J., Gulati, T., and Ganguly, K. (2019). Competing roles of slow oscillations and delta waves in memory consolidation versus forgetting. Cell, 179(2):514–526.

Kingma, D. P. (2014). Adam: A method for stochastic optimization. arXiv preprint arXiv:1412.6980.

Lawrence, N. (2003). Gaussian process latent variable models for visualisation of high dimensional data. Advances in neural information processing systems, 16.

Levenstein, D., Buzsáki, G., and Rinzel, J. (2019). Nrem sleep in the rodent neocortex and hippocampus reflects excitable dynamics. Nature communications, 10(1):2478.

Li, T. and Camera, G. L. (2025). A sticky poisson hidden markov model for solving the problem of over-segmentation and rapid state switching in cortical datasets. PloS one, 20(7):e0325979.

Liu, C., Todorova, R., Tang, W., Oliva, A., and Fernandez-Ruiz, A. (2023). Associative and predictive hippocampal codes support memory-guided behaviors. Science, 382(6668):eadi8237.

Liu, H., Ong, Y.-S., Shen, X., and Cai, J. (2020). When gaussian process meets big data: A review of scalable gps. IEEE transactions on neural networks and learning systems, 31(11):4405–4423.

Lopes-dos Santos, V., Ribeiro, S., and Tort, A. B. (2013). Detecting cell assemblies in large neuronal populations. Journal of neuroscience methods, 220(2):149–166.

Loren Frank Lab. track_linearization: 2d to 1d position linearization using hmms, (2024). URL https://github.com/LorenFrankLab/track_linearization.

Maaten, L. v. d. and Hinton, G. (2008). Visualizing data using t-sne. Journal of machine learning research, 9(Nov):2579–2605.

Maboudi, K., Ackermann, E., de Jong, L. W., Pfeiffer, B. E., Foster, D., Diba, K., and Kemere, C. (2018). Uncovering temporal structure in hippocampal output patterns. Elife, 7:e34467.

MacDonald, C. J., Lepage, K. Q., Eden, U. T., and Eichenbaum, H. (2011). Hippocampal “time cells” bridge the gap in memory for discontiguous events. Neuron, 71(4):737–749.

MacDonald, C. J., Carrow, S., Place, R., and Eichenbaum, H. (2013). Distinct hippocampal time cell sequences represent odor memories in immobilized rats. Journal of Neuroscience, 33(36):14607–14616.

Mallory, C. S., Widloski, J., and Foster, D. J. (2025). The time course and organization of hippocampal replay. Science, 387(6733):541–548.

Mazzucato, L., La Camera, G., and Fontanini, A. (2019). Expectation-induced modulation of metastable activity underlies faster coding of sensory stimuli. Nature neuroscience, 22(5):787–796.

McInnes, L., Healy, J., and Melville, J. (2018). Umap: Uniform manifold approximation and projection for dimension reduction. arXiv preprint arXiv:1802.03426.

Meyers, E. M., Freedman, D. J., Kreiman, G., Miller, E. K., and Poggio, T. (2008). Dynamic population coding of category information in inferior temporal and prefrontal cortex. Journal of neurophysiology, 100(3):1407–1419.

Miller, P. and Katz, D. B. (2010). Stochastic transitions between neural states in taste processing and decision-making. Journal of Neuroscience, 30(7):2559–2570.

Monaco, J. D., Rao, G., Roth, E. D., and Knierim, J. J. (2014). Attentive scanning behavior drives one-trial potentiation of hippocampal place fields. Nature neuroscience, 17(5):725–731.

O’Keefe, J. and Speakman, A. . (1987). Single unit activity in the rat hippocampus during a spatial memory task. Experimental brain research, 68(1):1–27.

Pandarinath, C., O’Shea, D. J., Collins, J., Jozefowicz, R., Stavisky, S. D., Kao, J. C., Trautmann, E. M., Kaufman, M. T., Ryu, S. I., Hochberg, L. R., et al. (2018). Inferring single-trial neural population dynamics using sequential auto-encoders. Nature methods, 15(10): 805–815.

Parks, D. F., Schneider, A. M., Xu, Y., Brunwasser, S. J., Funderburk, S., Thurber, D., Blanche, T., Dyer, E. L., Haussler, D., and Hengen, K. B. (2024). A nonoscillatory, millisecond-scale embedding of brain state provides insight into behavior. Nature neuroscience, 27(9):1829–1843.

Peyrache, A., Khamassi, M., Benchenane, K., Wiener, S. I., and Battaglia, F. P. (2009). Replay of rule-learning related neural patterns in the prefrontal cortex during sleep. Nature neuroscience, 12 (7):919–926.

Peyrache, A., Benchenane, K., Khamassi, M., Wiener, S. I., and Battaglia, F. P. (2010). Principal component analysis of ensemble recordings reveals cell assemblies at high temporal resolution. Journal of computational neuroscience, 29(1):309–325.

Pfeiffer, B. E. and Foster, D. J. (2013). Hippocampal place-cell sequences depict future paths to remembered goals. Nature, 497(7447):74–79.

Pillow, J. W., Shlens, J., Paninski, L., Sher, A., Litke, A. M., Chichilnisky, E., and Simoncelli, E. P. (2008). Spatio-temporal correlations and visual signalling in a complete neuronal population. Nature, 454(7207):995–999.

Pouget, A. and Snyder, L. H. (2000). Computational approaches to sensorimotor transformations. Nature neuroscience, 3(11):1192–1198.

Quian Quiroga, R. and Panzeri, S. (2009). Extracting information from neuronal populations: information theory and decoding approaches. Nature Reviews Neuroscience, 10(3):173–185.

Rabiner, L. R. (2002). A tutorial on hidden markov models and selected applications in speech recognition. Proceedings of the IEEE, 77(2): 257–286.

Ramanathan, D. S., Gulati, T., and Ganguly, K. (2015). Sleep-dependent reactivation of ensembles in motor cortex promotes skill consolidation. PLoS biology, 13(9):e1002263.

Rubin, A., Sheintuch, L., Brande-Eilat, N., Pinchasof, O., Rechavi, Y., Geva, N., and Ziv, Y. (2019). Revealing neural correlates of behavior without behavioral measurements. Nature communications, 10(1): 4745.

Saper, C. B., Fuller, P. M., Pedersen, N. P., Lu, J., and Scammell, T. E. (2010). Sleep state switching. Neuron, 68(6):1023–1042.

Sato, M., Mizuta, K., Islam, T., Kawano, M., Sekine, Y., Takekawa, T., Gomez-Dominguez, D., Schmidt, A., Wolf, F., Kim, K., et al. (2020). Distinct mechanisms of over-representation of landmarks and rewards in the hippocampus. Cell reports, 32(1).

Steinmetz, N. A., Zatka-Haas, P., Carandini, M., and Harris, K. D. (2019). Distributed coding of choice, action and engagement across the mouse brain. Nature, 576(7786):266–273.

Steriade, M., Nunez, A., and Amzica, F. (1993). A novel slow (< 1 hz) oscillation of neocortical neurons in vivo: depolarizing and hyperpolarizing components. Journal of neuroscience, 13(8):3252–3265.

Sun, W., Winnubst, J., Natrajan, M., Lai, C., Kajikawa, K., Bast, A., Michaelos, M., Gattoni, R., Stringer, C., Flickinger, D., et al. (2025). Learning produces an orthogonalized state machine in the hippocampus. Nature, 640(8057):165–175.

Takigawa, M., Huelin Gorriz, M., Tirole, M., and Bendor, D. (2024). Evaluating hippocampal replay without a ground truth. ELife, 13: e85635.

Tenenbaum, J. B., Silva, V. d., and Langford, J. C. (2000). A global geometric framework for nonlinear dimensionality reduction. science, 290(5500):2319–2323.

Tingley, D. and Buzsáki, G. (2020). Routing of hippocampal ripples to subcortical structures via the lateral septum. Neuron, 105(1):138–149.

Tingley, D. and Peyrache, A. (2020). On the methods for reactivation and replay analysis. Philosophical Transactions of the Royal Society B: Biological Sciences, 375(1799).

Van de Ven, G. M., Trouche, S., McNamara, C. G., Allen, K., and Dupret, D. (2016). Hippocampal offline reactivation consolidates recently formed cell assembly patterns during sharp wave-ripples. Neuron, 92(5):968–974.

van der Meer, M. A., Kemere, C., and Diba, K. (2020). Progress and issues in second-order analysis of hippocampal replay. Philosophical Transactions of the Royal Society B: Biological Sciences, 375(1799).

Vollan, A. Z., Gardner, R. J., Moser, M.-B., and Moser, E. I. (2025). Left–right-alternating theta sweeps in entorhinal–hippocampal maps of space. Nature, 639(8056):995–1005.

Watson, B. O., Levenstein, D., Greene, J. P., Gelinas, J. N., and Buzsáki, G. (2016). Network homeostasis and state dynamics of neocortical sleep. Neuron, 90(4):839–852.

Widloski, J. and Foster, D. J. (2022). Flexible rerouting of hippocampal replay sequences around changing barriers in the absence of global place field remapping. Neuron, 110(9):1547–1558.

Williams, A. H., Kim, T. H., Wang, F., Vyas, S., Ryu, S. I., Shenoy, K. V., Schnitzer, M., Kolda, T. G., and Ganguli, S. (2018). Unsupervised discovery of demixed, low-dimensional neural dynamics across multiple timescales through tensor component analysis. Neuron, 98 (6):1099–1115.

Williams, C. K. and Rasmussen, C. E. Gaussian processes for machine learning, volume 2. MIT press Cambridge, MA, (2006).

Wu, A., Roy, N. A., Keeley, S., and Pillow, J. W. (2017). Gaussian process based nonlinear latent structure discovery in multivariate spike train data. Advances in neural information processing systems, 30.

Yang, W., Sun, C., Huszár, R., Hainmueller, T., Kiselev, K., and Buzsáki, G. (2024). Selection of experience for memory by hippocampal sharp wave ripples. Science, 383(6690):1478–1483.

Yu, B. M., Cunningham, J. P., Santhanam, G., Ryu, S., Shenoy, K. V., and Sahani, M. (2008). Gaussian-process factor analysis for low-dimensional single-trial analysis of neural population activity. Advances in neural information processing systems, 21.

Zhang, K., Ginzburg, I., McNaughton, B. L., and Sejnowski, T. J. (1998). Interpreting neuronal population activity by reconstruction: unified framework with application to hippocampal place cells. Journal of neurophysiology, 79(2):1017–1044.

Zheng, C., Bieri, K. W., Hsiao, Y.-T., and Colgin, L. L. (2016). Spatial sequence coding differs during slow and fast gamma rhythms in the hippocampus. Neuron, 89(2):398–408.

Zheng, Z. S., Huszár, R., Hainmueller, T., Bartos, M., Williams, A. H., and Buzsáki, G. (2024). Perpetual step-like restructuring of hippocampal circuit dynamics. Cell reports, 43(9).

Zhou, D. and Wei, X.-X. (2020). Learning identifiable and interpretable latent models of high-dimensional neural activity using pi-vae. Advances in neural information processing systems, 33:7234–7247.

Zutshi, I., Apostolelli, A., Yang, W., Zheng, Z. S., Dohi, T., Balzani, E., Williams, A. H., Savin, C., and Buzsáki, G. (2025). Hippocampal neuronal activity is aligned with action plans. Nature, 639(8053): 153–161.

